# PIEZO2‐dependent rapid pain system in humans and mice

**DOI:** 10.1101/2023.12.01.569650

**Authors:** Otmane Bouchatta, Marek Brodzki, Houria Manouze, Gabriela B. Carballo, Emma Kindström, Felipe M. de‐Faria, Huasheng Yu, Anika R. Kao, Oumie Thorell, Jaquette Liljencrantz, Kevin K. W. Ng, Eleni Frangos, Bengt Ragnemalm, Dimah Saade, Diana Bharucha‐Goebel, Ilona Szczot, Warren Moore, Katarzyna Terejko, Jonathan Cole, Carsten Bonnemann, Wenquin Luo, David A. Mahns, Max Larsson, Gregory J. Gerling, Andrew G. Marshall, Alexander T. Chesler, Håkan Olausson, Saad S. Nagi, Marcin Szczot

**Affiliations:** Center for Social and Affective Neuroscience, Linköping University, Linköping, Sweden; Department of Neuroscience, Perelman School of Medicine, University of Pennsylvania, Philadelphia, USA; School of Engineering and Applied Science, University of Virginia, Charlottesville, USA; School of Medicine, Western Sydney University, Sydney, Australia; National Center for Complementary and Integrative Health, National Institutes of Health, Bethesda, USA; Department of Anesthesiology and Intensive Care, Institute of Clinical Sciences, Sahlgrenska Academy at University of Gothenburg, Gothenburg, Sweden; National Institute of Neurological Disorders and Stroke, National Institutes of Health, Bethesda, USA; Institute of Life Course and Medical Sciences, University of Liverpool, UK; Biology of Astrocytes Research Group, Łukasiewicz Research Network ‐ PORT Polish Center for Technology Development, Wroclaw, Poland; University Hospitals, Dorset, and University of Bournemouth, UK

## Abstract

The PIEZO2 ion channel is critical for transducing light touch into neural signals but is not considered necessary for transducing acute pain in humans. Here, we discovered an exception – a form of mechanical pain evoked by hair pulling. Based on observations in a rare group of individuals with PIEZO2 deficiency syndrome, we demonstrated that hair-pull pain is dependent on PIEZO2 transduction. Studies in control participants showed that hair-pull pain triggered a distinct nocifensive response, including a nociceptive reflex. Observations in rare Aβ deafferented individuals and nerve conduction block studies in control participants revealed that hair-pull pain perception is dependent on Aβ input. Single-unit axonal recordings revealed that a class of cooling-responsive myelinated nociceptors in human skin is selectively tuned to painful hair-pull stimuli. Further, we pharmacologically mapped these nociceptors to a specific transcriptomic class. Finally, using functional imaging in mice, we demonstrated that in a homologous nociceptor, Piezo2 is necessary for high-sensitivity, robust activation by hair-pull stimuli. Together, we have demonstrated that hair-pulling evokes a distinct type of pain with conserved behavioral, neural, and molecular features across humans and mice.

## Introduction

Sensing environmental threats is an essential role of the somatosensory system. In recent decades, our understanding of how the peripheral nervous system differentiates between changes in temperature or submodalities of touch has broadened. However, despite people’s ability to discern various types of noxious mechanical stimuli, such as a pinprick or a pinch, the neural mechanisms underlying this capability remain less understood. Traditionally, human sensory neurons are categorized based on unique physiological properties^1^. Textbook distinctions rely on the dichotomy between fast ‘first pain’, mediated by thinly myelinated (Aδ) nociceptors, and slow ‘second pain’, mediated by unmyelinated (C) nociceptors, whereas tactile modalities are conveyed by thickly myelinated ultrafast (Aβ) low-threshold receptors^2^. However, we have shown that a hitherto unknown population of Aβ brush-insensitive mechanoreceptors with high indentation thresholds (Aβ mechanonociceptors) contributes to acute pricking pain in humans, challenging the conventional view that pain signaling is slower than touch signaling^3^. Furthermore, a subset of these Aβ nociceptors responds to cooling^4^, illustrating that the functional diversity of nociceptors may be greater than previously appreciated.

A conventional clinical pain assessment tool, such as the McGill pain questionnaire^2^, contains multiple qualitative categories and descriptors for mechanical pain. However, little is known about the cellular basis of the different submodalities of mechanical pain, such as punctate, constrictive, or traction pain, associated with descriptors like pricking, pinching, or tugging. Recent advances in single-cell RNA sequencing have opened up new possibilities for delineating the molecular makeup of human sensory neurons^4-7^. In particular, single-soma deep RNA-sequencing of human DRG neurons has revealed several populations of putative mechanonociceptors outlined by the expression of the *SCN10A* gene^4^, indicating a larger diversity of mechanonociceptors than identified so far functionally^3,8-10^. Therefore, to identify the cellular basis of mechanical pain diversity, a physiological characterization needs to be matched with molecular identification.

Many classes of *SCN10A*^*+*^ neurons express the mechanotransduction channel PIEZO2^11^. This channel serves as a primary transducer for touch sensation in both mice and humans^12,13^, and it has a critical role in damage- or disease-evoked forms of pathophysiological pain, e.g., tactile allodynia^14,15^ and visceral pain hypersensitivity^16,17^. PIEZO2 has been implicated in contributing to transduction of some types of acute pain in mice^14,18,19^ but is not known to be required for any distinct pain submodality^13,15^. Further, individuals with PIEZO2-deficiency syndrome retain pain sensation despite their inability to perceive discriminative touch^3,13,15^.

In the current study, examination of PIEZO2-deficiency syndrome individuals has revealed a deficit in hair-pull pain perception. Using targeted electrophysiological and psychophysical approaches in control participants, including single-fiber recordings, we have identified hair-pull pain as a distinct Aβ-dependent submodality of mechanical pain, encoded by a class of brush-insensitive mechanoreceptors in the skin. Furthermore, using a pharmacological mapping approach, we have confirmed that these hair-pull coding fibers belonged exclusively to a transcriptomically defined class of nociceptors. Moreover, using a conditional knockout of Piezo2 in the mouse homologs of hair-follicle nociceptors, we have shown that Piezo2 is critical for hair-pulling response of these neurons. Our data demonstrate that a specialized class of rapidly conducting PIEZO2-dependent nociceptors associated with hair follicles serves as evolutionarily conserved mediators of hair-pull pain.

## Results

### Hair-pulling pain is a distinct PIEZO2-dependent mechanical pain submodality mediated by rapidly conducting fibers

We have previously identified a rare group of individuals carrying inactivating variants in the PIEZO2 gene with compound heterozygosity^13,15^. While individuals with PIEZO2-deficiency syndrome have difficulty producing coordinated movements and exhibit profound touch deficits, their ability to perceive mechanical pain evoked by indentation, pinching, and pressure is preserved^3,13,15^. We now report that these individuals exhibit a profound pain deficit in response to the pulling of single hairs (Figure 1A, B). In a psychophysical characterization of scalp hair-pull pain sensation, control participants consistently reported hair-pulling as painful, while individuals with PIEZO2-deficiency syndrome, when subjected to the same pull forces, predominantly reported them as nonpainful (Figure 1A, B). All subsequent human experiments involved control participants. We assessed if noxious pulling of single hairs evoked a distinct percept compared to other forms of mechanical pain. When participants were presented with single hair-pull or pinprick stimuli at equivalent pain intensities, they could reliably distinguish between them (Figure 1C). Pain quality data revealed a mix of common descriptors (such as ‘sharp’) and those that were preferential to pinprick (‘stabbing’) and hair-pull (‘hot-burning’ and ‘tender’, Extended Figure 1). Notably, the stimulation force required to achieve similar pain intensity ratings was many times lower for hair pull compared to pinprick stimulation. To investigate this further, we measured four skin deformation metrics (see Methods and Extended Figure 2) during single-hair pulling and skin indentation. Despite equivalent skin deformation between hair pulling and indentation, we observed that pain ratings were still higher during hair-pull than during indentation (Extended Figure 2).

**Figure 1:**
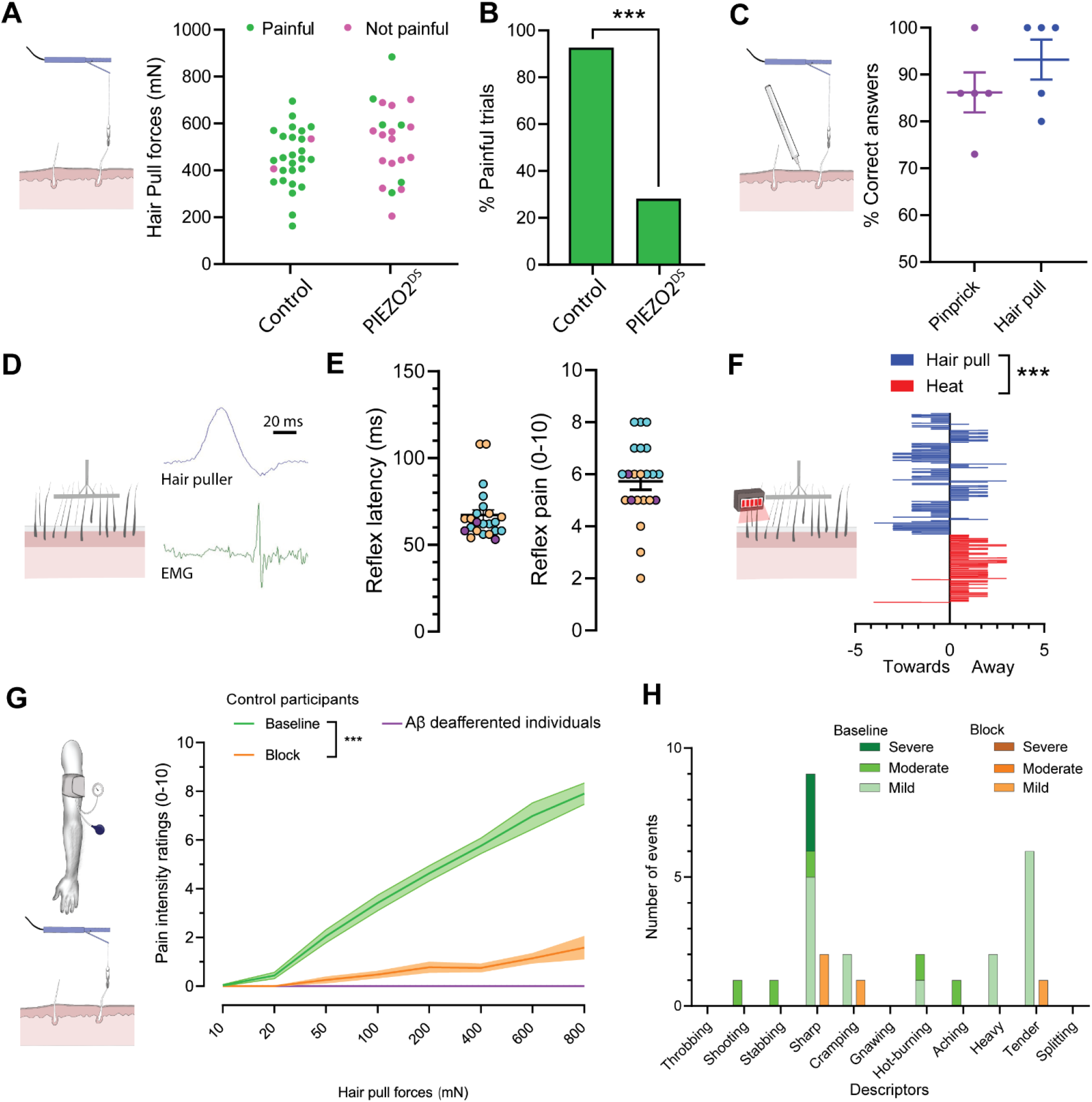
Hair-pulling pain is a distinct PIEZO2-dependent pain submodality mediated by rapidly conducting fibers. **A-B**. Individuals with PIEZO2 deficiency syndrome (PIEZO2^DS^) exhibited a profound deficit in hair-pull pain perception (control participants: n=7, PIEZO2^DS^ individuals: n=6). **A**. At comparable forces, most hair-pull trials were reported as painful by control participants and nonpainful by PIEZO2^DS^ individuals. Dots represent individual trials (control participants: n=28 trials, PIEZO2^DS^ individuals: n=21 trials). **B**. The histograms represent the percentage of trials reported as painful (control participants: 92.8% *vs*. PIEZO2^DS^ individuals: 28.57%, p<0.0001; Fisher’s exact test). **C**. Hair-pulling evoked a readily discriminable percept. Control participants consistently distinguished between hair-pull pain and pinprick pain at equivalent pain intensity ratings (correct responses: hair pull 93.2 ± 6.0%, pinprick 86.2 ± 7.1%; n=15 trials per stimulus per participant; n=5 participants). Data are presented as individual- and pooled-mean (± SEM) responses in percentages. D-E. Hair-pull pain elicited a nociceptive reflex. **D**. An example EMG recording of a nociceptive reflex response evoked by multi-hair pulling from a posterior arm muscle. **E**. Latency distribution of reflex responses recorded in anterior and posterior muscle compartments of the upper arm and corresponding pain ratings. Since no differences were found in reflex latencies and pain ratings (p>0.05; t-test) between the two compartments, the data were pooled. All reflex responses (n=23) were evoked at stimulus intensities rated as painful. Data are shown as individual and mean (±SEM) responses from 3 control participants (color-coded). **F**. Hair pulling evoked a distinct urge-to-move psychophysical response. Control participants chose to preferentially move towards the pain source in response to multi-hair pulling contrary to heating where they chose to move away (n=11 participants, p<0.001; Mann Whitney test, with hair pull urge *vs*. baseline: p=0.0585 and heat urge *vs*. baseline: p<0.001; Paired t-test). Data are presented as mean ± SEM. **G-H**. Hair-pull pain was scalable and mediated by Aβ fibers. **G**. Pain intensity ratings in response to single-hair pulling forces were assessed in control participants (n=20) at ‘Baseline’ and during preferential Aβ-fiber conduction ‘Block’, as well as in individuals with selective Aβ deafferentation (n=2). Pain intensity ratings were significantly affected by hair-pulling forces (F_(7,438)_ = 13.50, p<0.0001) and whether large myelinated fibers were conducting or not (F_(2,438)_ = 266.5, p<0.0001). During the conduction block, the hair-pull pain ratings decreased significantly across a range of pull forces (Baseline *vs*. Block: [50-800 mN]: p<0.0001). In addition, the Aβ-deafferented individuals were unable to perceive hair-pull pain. Statistical differences were assessed using a two-way ANOVA followed by Tukey’s multiple comparisons test. Data are presented as mean ± SEM. **H**. Quality of hair-pull pain assessed using the short-form McGill questionnaire in Baseline and Block conditions. The data show the number of occurrences (events) for each pain descriptor.

**Figure 2:**
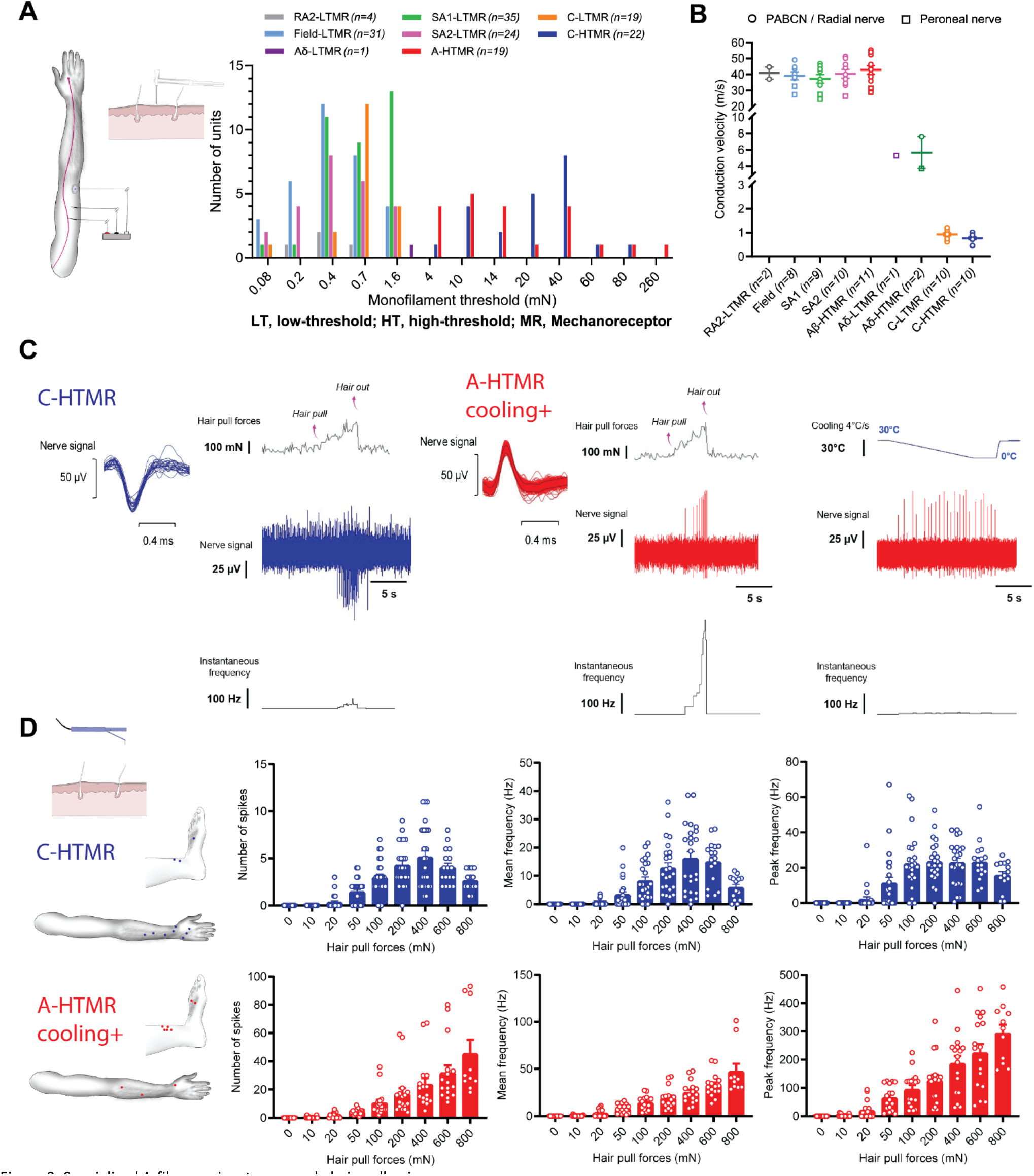
Specialized A-fiber nociceptors encode hair-pull pain. **A**. Mechanical threshold distribution of HTMRs and LTMRs in the recorded sample. For RA1 afferents, the preferred stimulus is hair movement, so monofilament thresholds were not measured. **B**. Conduction velocities of HTMRs and LTMRs in response to surface electrical stimulation and/or mechanical stimulation using an electronic monofilament with force-feedback. Conduction velocities of Aβ-HTMRs (42.8 ± 8.9 m/s, n=11) were statistically indistinguishable from Aβ-LTMRs ([SA1]: 37.2 ± 7.8 m/s, q=2.74, p=0.5318, n=9; [SA2]: 40.4 ± 8.3 m/s, q=1.21, p=0.9890, n=10; [Field]: 39.2 ± 6.9 m/s, q=1.726, p=0.922, n=8). Statistical differences were assessed using a one-way ANOVA with Tukey’s multiple comparisons test. Data are shown as individual and mean (±SEM) responses. PABCN, posterior antebrachial cutaneous nerve. **C**. Representative traces displaying hair-pull responses of a C-HTMR and a cooling-responsive A-HTMR. Single hairs were pulled from the receptive field of the recorded afferent until the hair was extracted. **D**. Receptive field locations of C-HTMRs (n=11) and cooling-responsive A-HTMRs (n=10) with hair present in the receptive field for hair-pull testing (left). The data show the individual trials and mean (±SEM) responses of C-HTMRs and cooling+ A-HTMRs to hair pulling at different forces. For C-HTMRs, hair pulling forces had a significant effect on the number of spikes (F_(8,190)_ = 29.36, p<0.0001), as well as on mean (F_(8,190)_ = 20.21, p<0.0001) and peak frequencies (F_(8,190)_ = 20.81, p<0.0001). The response of C-HTMRs to increasing pull forces plateaued or dropped with further increases in pull forces. Accordingly, the peak frequency at the highest pulling force of 800 mN was not significantly different from other forces tested (p>0.05). For cooling+ A-HTMRs, hair pulling forces also had a significant effect on the number of spikes (F_(8,135)_ = 17.39, p<0.0001), as well as on mean (F_(8,135)_ = 31.34, p<0.0001) and peak frequencies (F_(8,135)_ = 28.19, p<0.0001). However, in contrast to C-HTMRs, the activity in A-HTMRs was tuned to increasing hair-pull forces with the most robust response observed at the highest pulling force. Accordingly, the peak frequency at the highest pulling force of 800 mN was significantly higher than all other forces tested (at least p<0.001). Statistical differences were assessed by a one-way ANOVA with Tukey’s multiple comparisons test. The mechanical threshold, conduction velocity (where tested), and cooling response of five A-HTMRs from this sample were published in a preprint^4^.

Next, we explored whether hair-pull pain triggers a nociceptive reflex. To achieve this, we developed an automated multi-hair puller with force feedback. We observed reproducible, short-latency (67.1 ± 3.1 ms) EMG responses in upper arm muscles in response to painful hair-pull stimuli delivered to the forearm (Figure 1D, E). No reflex responses were observed below the pain threshold (Figure 1E). Acute pain helps one avoid potentially harmful stimuli; therefore, the typical reaction involves a withdrawal response (e.g. Thorell et al.^20^). However, when hair is pulled, the appropriate protective behavior would be to approach towards the source of the stimulus rather than move away from it. To explore this further, we conducted a psychophysical task in which participants were provided with an urge-to-move scale with endpoints ‘towards’ and ‘away’ and were asked to rate their preferred response during painful multi-hair pulling or heat-induced pain. We observed that hair-pulling preferentially induced an urge to approach the stimulus, contrary to heating, which evoked an urge to withdraw from the stimulus (Figure 1F), showing that hair-pull pain elicits a distinct nocifensive behavior. After establishing hair-pull as a distinct type of mechanical pain, we sought to characterize the primary afferent class necessary for mediating this submodality. First, we tested a range of pulling forces applied to single hairs in the forearm, hand, and foot regions. Since no differences (p>0.05) were found in pain ratings between these regions, the data were pooled for subsequent analysis. Importantly, the pain ratings increased with the pulling force, indicating that hair-pull pain was a scalable sensation (Figure 1G). Then, we used a progressive nerve ischemic block, induced by a blood pressure cuff^20^, to dissociate the contributions of different afferent fiber classes to hair-pull pain perception. The progression of the block was tracked using quantitative sensory evaluations with vibration intensity and cooling and warming detection taken as measures for Aβ-, Aδ-, and C-fiber function, respectively^21^. Under conditions where the sense of vibration intensity was almost abolished (Extended Figure 3A), while the thermal stimuli could still be detected within the nonpainful range (Extended Figures 3B-C), we found that the perception of hair-pull pain was abolished (Figure 1G). This effect was also reflected in the pain quality reports with a reduced frequency of chosen descriptors, most notably, ‘sharp’ and ‘tender’, during the block (Figure 1H). Nerve blocks, while effective, rely on functional readouts, and it cannot be ruled whether any residual large-fiber afferents were still conducting; further, during the block, the thermal sensibility was also reduced. Therefore, we tested hair-pull pain perception in two well-characterized patients with a rare, selective Aβ deafferentation^22^ and found that neither of them perceived hair-pulling stimuli as painful, demonstrating the critical role of very rapidly conducting fibers in hair-pull nociception (Figure 1G).

**Figure 3:**
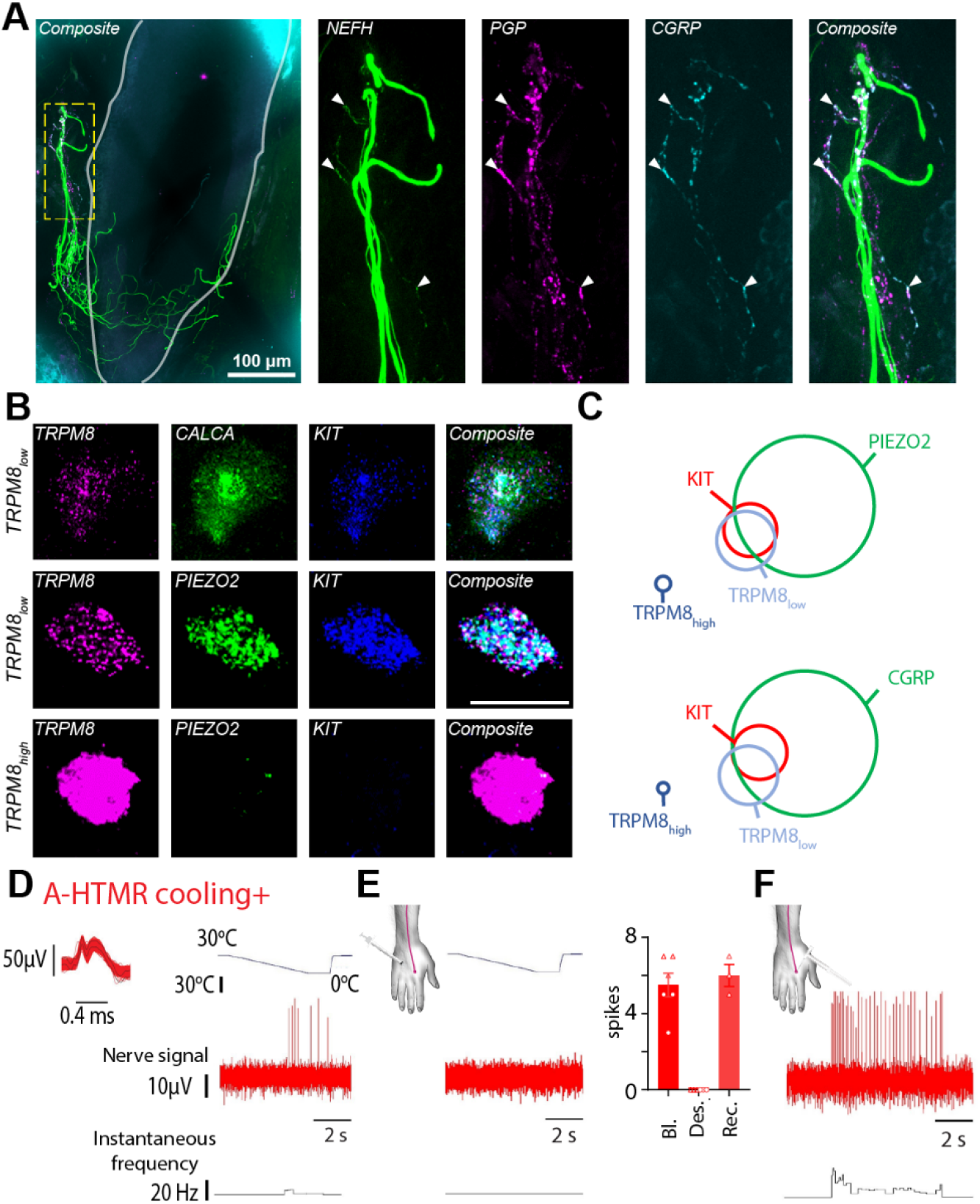
Hair follicle-associated myelinated nociceptors belonged to a distinct transcriptomic class. **A**. Human skin immunofluorescence showed colocalization of Nefh, PGP and CGRP markers in nerve endings associated with hair follicles, confirming the existence of myelinated hair-follicle nociceptors in humans. Solid grey line marks the hair follicle. **B**. Fluorescent in situ hybridization (FISH) in human DRG neurons showed co-expression of TRPM8^low^ with genetic markers delineating the hair-follicle nociceptor population. Scale bar = 50 μm. **C**. Quantification of FISH data (n=180 cells from 2 donors for KIT-PIEZO2-TRPM8 and 204 cells from 2 donors for KIT-CGRP-TRPM8). KIT-PIEZO2-TRPM8: KIT 8.3%, PIEZO2 40.6%, TRPM8^low^ 12.2%, TRPM8^high^ 3.3% of all neurons. KIT-CALCA-TRPM8: KIT 6.4%, CALCA 45.6%, TRPM8^low^ 10.8%, TRPM8^high^ 2.0% of all neurons. **D-F**. Cooling responsiveness of cooling+ A-fiber nociceptors is dependent on TRPM8. Recording traces showing the response of a cooling+ A-fiber nociceptor to a drop in temperature before **(D)** and after menthol injections **(E)**, inducing TRPM8 desensitization. Individual and mean (±SEM) responses of cooling+ A-fiber nociceptors to a drop in temperature under ‘Baseline (Bl)’ and ‘TRPM8 desensitization (Des.)’ conditions (2 units, tested in triplicate). In one case, the ‘Recovery (Rec.)’ phase was also recorded. **F**. Recording trace showing the response of a cooling+ A-fiber nociceptor to high-threshold mechanical stimulation (600 mN von Frey) under ‘TRPM8 desensitization’ condition.

Taken together, we have identified hair-pulling pain as a distinct mechanical pain submodality. This type of pain is characterized by unique psychophysical and molecular properties, the hallmark of which is a high sensitivity, A-fiber dependence, and PIEZO2 reliance.

### Specialized Aβ-HTMRs encode hair-pull sensation in the noxious range

Psychophysics helps us describe reactions and perceptions triggered by painful stimuli but cannot identify the underlying physiological mechanisms. Therefore, to test what type of neurons are involved in the neural coding of single hair pull, we used the microneurography technique. Microneurography affords the unique ability to record the activity of single peripheral afferents in awake human participants^23^. Importantly, single-unit activity can be used to map tuning properties and receptive fields and correlate these quantitative measures with a person’s subjective experience. In the current study, we performed recordings from the radial, antebrachial, and superficial peroneal nerves of 94 control participants (Figure 2A, Extended Figure 4). We recorded from a total of 180 cutaneous afferents (peroneal, 30; radial, 132; antebrachial, 18), including multiple brush-sensitive types: Aβ field-low threshold mechanoreceptor (LTMR) (n = 31), Aβ rapidly adapting type 1 (RA1) LTMR (hair unit, n = 25), Aβ RA2-LTMR (Pacinian unit, n = 4), Aβ slowly adapting type 1 (SA1) LTMR (n = 35), Aβ SA2-LTMR (n = 24), Aδ-LTMR (n = 1), and C-LTMR (n = 19). All brush-sensitive afferents had low mechanical thresholds to skin indentation (<4 mN; Figure 2A). We also recorded from brush-insensitive types, comprising A-fiber high-threshold mechanoreceptor (HTMR) (n = 19) and C-HTMR (n = 22). The brush-insensitive afferents had high mechanical thresholds to indentation (≥4 mN; Figure 2A), and all responded to pinching. The recorded afferents had conduction velocities (where tested) in the Aβ- (>30 m/s) or C-fiber (<2 m/s) range, barring two HTMRs and an LTMR conducting in the slow Aδ range (Figure 2B). The receptive field locations of all recorded afferents are shown in Extended Figure 4.

**Figure 4:**
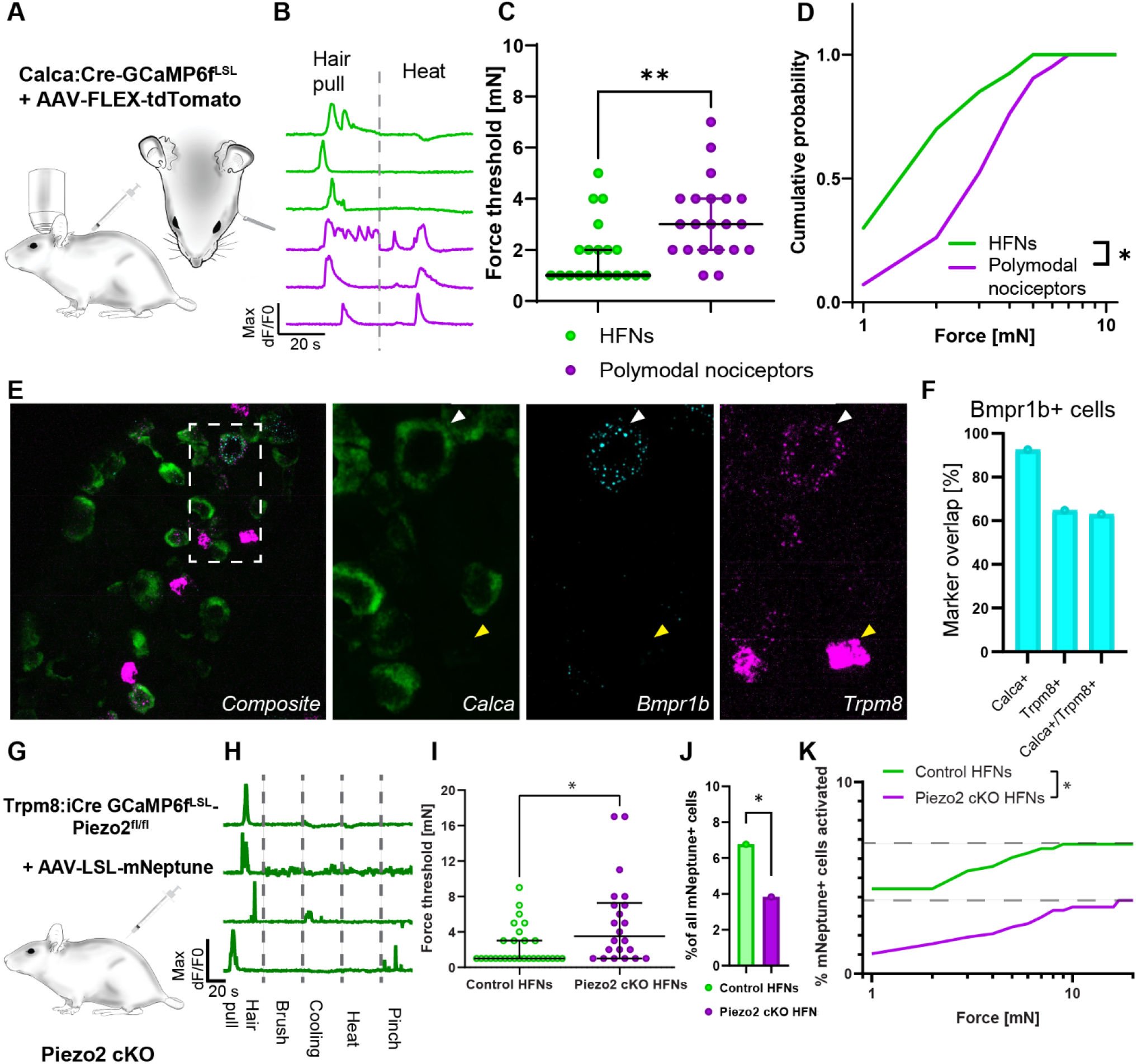
Piezo2 set the sensitivity of hair follicle nociceptors to hair pulling. **A**. Diagram of the trigeminal in vivo calcium imaging experiment with single hair-pull stimuli and genetic model for labelling *Calca* nociceptors. *Calca*^*Cre*^ mice were crossed with Ai95 Cre dependent GCamp6f strain. To postnatally limit the labelling to persistent Calca nociceptors offspring was injected at P1-P3 with AAV/PHP.S-CAG-Fl-tdTomato virus. **B**. Representative calcium transients recorded in *Calca*^*+*^ heat-insensitive hair follicle nociceptors (green) and heat-responsive polymodal nociceptors (magenta) neurons evoked by single hair-pull and heat stimuli. **C**. Forces activating HFNs (green) were markedly smaller (median threshold: 1 mN, n=20) than for polymodal nociceptors (magenta, median threshold: 3 mN, p=0.0037, n=21 for polymodal nociceptors, unpaired Student’s t-test). Data is shown as individual replicates with median and interquartile range overlay. **D**. Cumulative distribution of activation thresholds for HFNs (green) and polymodal nociceptors (magenta) neurons evoked by single hair-pull shows a markedly higher sensitivity to hair-pull in HFNs. Statistical differences were assessed by two-way ANOVA (F _(1, 10)_=6.521, p=0.0287). **E**. Mouse HFNs express low levels of *Trpm8*. Sample images of fluorescence in situ hybridization co-localization of HFN’s markers *Bmrp1b, Calca* and *Trpm8*. **F**. *Bmpr1b*^*+*^ neurons express Calca and majority of them expresses low levels of *Trpm8* transcripts. **G**. Genetic model of Piezo2 loss of function in HFNs. GCamp6f expressing conditional Piezo2 KO was generated by crossing the Trpm8^iCre^ mouse strain with double transgenic Ai95-Piezo2^fl/fl^ line to obtain homozygous Piezo2 cKO. To limit the labelling to persistent Calca nociceptors offspring was injected at P1-P3 with AAV/PHP.S-CAG-LSL-mNeptune. **H**. Representative calcium transients of functionally isolated HFNs recorded in control mice. Demonstrating high selectivity of HFNs. **I**. Hair follicle nociceptors (green) sensitivity to hair pull force (median threshold: 1 mN, n=29) was markedly diminished in the absence of Piezo2 (magenta, median threshold: 3.5 mN, n=21, p=0.0158, upaired Student’s t-test). Data is shown as individual replicates with median and interquartile range overlay. **J**. Percentage of cells responding to hair-pull in Piezo2cKO was diminished by almost half (3.8% in Piezo2 cKO vs 6.8% in control, p=0.0418, Fisher’s exact test). **K**. Additive effect of Piezo2 on hair follicle nociceptor responses during hair-pull. Cumulative distribution of hair follicle nociceptors activation scaled to total recruitment during hair-pull trial, (two-way ANOVA (F_(1, 19)_=3556, p<0.0001, n=29 neurons from 7 mice for control and n=22 neurons from 9 mice for Piezo2KO).

The fast-conducting A-HTMRs can be categorized into two types in humans: both types respond to noxious skin indentations, but only one type responds to cooling ^4^. Neither type responds to heating. In the current study, we uncovered a novel association between cooling and hair-pull responsiveness. While the cooling-unresponsive A-HTMRs did not respond to single-hair pulling (tested in 7 units; Extended Figure 6D), the cooling-responsive A-HTMRs showed selective tuning to painful pulling of single hairs within their receptive fields, manifesting as an increased response to progressively painful hair-pull forces (Figures 2C-D). That these A-HTMRs not only exhibit high thresholds as the name implies but also encode noxious stimuli classifies them as nociceptors. On the other hand, the C-HTMRs reached a plateau, and eventually, their response dropped off within the painful hair-pull range (Figures 2C-D). None of the LTMR classes displayed tuning in the painful range of hair-pull forces (Extended Figures 5 and 6), except the SA2-LTMRs, which exhibited a wide range of sensitivity from innocuous well into the noxious range of pull forces before they too eventually plateaued (Extended Figure 6A). That cooling+ A-fiber nociceptors were the only class that showed selective tuning to painful hair-pull forces suggests they are ideally suited for encoding hair-pull pain in humans.

### Hair follicle-associated myelinated nociceptors belong to a distinct transcriptomic class

The unique physiological signature of the A-fiber nociceptors that responded to hair pull and cooling suggests that these neurons might belong to a distinct molecular class. First, we aimed to establish that myelinated nociceptors are indeed associated with hair follicles. Using immunohistochemistry in human skin biopsies, we found that a subset of neurofilament heavy chain (NEFH) immunoreactive endings surrounding the hair follicle were also positive for the nociceptive marker calcitonin gene-related peptide (CGRP, encoded by CALCA, Figure 3A). This observation suggested the presence of a myelinated nociceptive nerve ending associated with the hair follicle. Second, based on recently established single-cell RNA sequencing (scRNAseq) cellular atlases of human DRG neurons, we wanted to identify which specific transcriptomic class these neurons belong to. Recent scRNAseq studies^4-7^ have revealed multiple transcriptomic classes of myelinated CALCA^+^ neurons. Moreover, in specific differentiation protocols, human iPSC-derived population of *in vitro* mechanoreceptors co-express TRPM8 and PIEZO2^24^. We hypothesized that *in vivo*, since cooling+ A-fiber nociceptors were electrophysiologically characterized by cooling responses, they might express the cooling- and menthol-activated ion channel TRPM8. Indeed, fluorescence in situ hybridization revealed that, in addition to the well-characterized cooling neurons expressing high levels of TRPM8 (TRPM8^high^), humans possessed a distinct subtype of TRPM8^low^ neurons that expressed PIEZO2, KIT, and CALCA (Figure 3B). This transcriptional profile of human TRPM8^low^ neurons matched a specific population of NEFH^+^ neurons identified in scRNAseq of human sensory neurons^4-6^. This data makes human TRPM8^low^ neurons a likely candidate for hair-follicle nociceptors. To further test this, we compared the cooling responsiveness before and after acute desensitization of peripheral TRPM8 channels in two cooling+ A-fiber nociceptors (Figure 3D and E). This was achieved through sequential intradermal injections of menthol around the receptive field of the recorded afferent^25^. Following desensitization (confirmed by the loss of sensation to menthol and diminished cold perception), we observed a complete loss of the cooling response from both recorded afferents (tested in triplicate; Figures 3E), while they still responded to skin indentation (Figures 3F). This suggests that the cooling+ A-fiber nociceptors rely on functional TRPM8 channels for their cooling sensitivity and, therefore, likely belong to the TRPM8^low^ class. These data strongly support the idea that TRPM8^low^ neurons are hair-follicle nociceptors (HFNs).

### Piezo2 sets the mechanical sensitivity of the hair-follicle nociceptors

Human microneurography experiments require multiple recording sessions to record from a single HFN, and there is a limited number of individuals with PIEZO2-deficiency syndrome available for testing. Therefore, we decided to test the hair follicle nociceptors’ transduction mechanism using an animal model. We have recently demonstrated that in mice, a class of *Calca*^*+*^*Trpv1*^*-*^ nociceptors consists of mouse A-HTMRs forming a unique type of circumferential nerve ending around the hair follicle^26^. Therefore, we set out to test whether these neurons are functionally and molecularly analogous to human HFNs. To administer a controlled single hair-pull in mice, we used force-feedback forceps to pull on a thread attached to a single guard hair, allowing us to record both the neuronal response to hair-pulling and its force threshold (Figure 4A). Neural activity was assayed using high-throughput calcium imaging *in vivo* of the trigeminal ganglia^26,27^.

To selectively record from mouse HFNs, we used transgenic Calca:Cre driver crossed to GCAMP6f^LSL^ reporter strain. Offsprings were postnatally labeled with adenoassociated virus (AAV) conditionally expressing tdTomato (Fig. 4A). This approach allowed us to express a calcium indicator and use red reporter expression to identify *Calca*^*+*^ nociceptors in our recordings. To further distinguish between *Calca*^*+*^*Trpv1*^*-*^ HFNs and *Calca*^*+*^*Trpv1*^*+*^ polymodal nociceptors, we classified hair-pulling responsive neurons based on their lack of response to noxious heating (50°C; Figure 4B). When we measured the forces required for the hair-pulling responses, we noted that the HFNs were three times more sensitive than polymodal nociceptors (median activation threshold: 1 mN vs 3 mN; Figure 4C, D). These observations showed that mouse HFNs recapitulated two functional hallmarks of human hair follicle nociceptors: high sensitivity to hair-pulling and the lack of heating response.

In mice, *Calca*^*+*^*Trpv1*^*-*^ HFNs are characterized by the expression of a single marker gene, *Bmpr1b*^*28*^. High-sensitivity fluorescence in situ hybridization (FISH) revealed that the majority of *Bmpr1b* neurons expressed low levels of *Trpm8*, similar to human HFNs (Figure 4E, F). When we further characterized the expression of *Trpm8* in mouse DRG, we observed that, in addition to the well-known cold-sensing high *Trpm8*-expressing neurons^29,30^, some neurons expressed low levels of *Trpm8* transcript and co-expressed *Scn10a* and *Piezo2* (Extended Figure 7A-C). *Trpm8*^*low*^ cells were *Calca*^*+*^, and about half of them expressed *Bmpr1b* (47%; Fig. 4E,F), whereas the remaining subset expressed *Trpv1* (Extended Figure 7D). These data indicated that in mice, the *Trpm8*^*low*^ labels a population of sensory neurons which is highly enriched in HFNs. To genetically manipulate HFNs, we used a previously unpublished transgenic strain of *Trpm8:iCre* mice (SMOC repository number: NM-KI-225064). We validated it as faithfully targeting *Trpm8*^low^ cells (Extended Figure 8A, B). When the *Trpm8:iCre* animals were injected with an AAV encoding Cre-dependent reporter we observed recombination in the CGRP-positive circumferential endings around hair follicles (Extended figure 8C, D). Next, we used *Trpm8:iCre* to generate conditional Piezo2 knockout and expressed GCaMP6f in *Trpm8:iCre* lineage cells. *Trpm8* is expressed by some neurons only in the prenatal stage^31^, therefore a transgenic cross can encompass a larger population that is characterized by expression analysis in adults. To circumvent this problem, we additionally injected an AAV encoding Cre-dependent red fluorescent reporter to restrict our analysis to *Trpm8* persistent cells (Figure 4G). As expected, in control animals, a large fraction (26%) of responding labelled neurons were sensitive to hair-pull and showed high degree of selectivity to this stimulus (Figure 4H, Extended Figure 9A, C). Remaining labelled neurons comprised of polymodal HTMRs and thermoreceptors; however, in line with transcriptomic prediction, we observed no LTMRs (Extended Figure 9A-B). Measurements of activation threshold again confirmed high sensitivity of HFNs to hair-pulling, and in the absence of Piezo2, the activation threshold increased almost threefold (Figure 4H). In hair/pulling experiments, the delivered force was always limited to the force needed for the hair to be plucked out. We noticed that for some neurons in Piezo2 cKO, threshold values were close to or even above the average force needed to pluck out the hair (11 mN, n=78 trials across 8 animals). Therefore, we hypothesized that even higher response thresholds might be masked by the fact that we did not observe responsive neurons because the hair was plucked out before activation occurred. Congruent with this prediction, we observed that the average number of cells responding to hair pulling was decreased by half (Figure 4I). Effects shown in Figure 4I and 4J are additive, which means that more Piezo2^+^-HFNs activate in response to even the most miniscule force (1 mN) than Piezo2 cKO HFNs to the whole act of pulling out the hair (Figure 4K). Therefore, the loss of Piezo2 in HFNs leads to a strong suppression of their activation during the physiological process of hair-pulling.

## Discussion

Functional diversity of nociceptors has been studied for over 50 years^32^ and while conduction speed is an important metric in clinical assessments of nerve function, the pursuit of precision treatments requires a deeper understanding of the molecular and cellular mechanisms underlying distinct pain submodalities. The conventional view attributes the pain sensation to nociceptors detecting thermal, chemical, or mechanical stimuli, or their combinations. However, the molecular genetics of human nociceptors reveal far greater diversity than can be ascribed based on functional characterization alone^33^, prompting further exploration into the molecular basis of diverse pain submodalities. In the current study, we have found that hair-follicle nociception encompasses functional, molecular, morphological, and cellular specialization, giving rise to a particular submodality of mechanical pain. Hairy skin covers most of our body, and the density of hair follicles is on par with that of tactile afferent endings^34-36^. Moreover, a complex sensory organ forms around each hair follicle, innervated by multiple classes of LTMRs^37-39^. In contrast, the nociceptive system protecting human hair follicles has, until now, remained largely unexplored despite the familiar experience of pain caused by hair removal.

Human skin is equipped with very fast-conducting nociceptors, which contribute to the pinprick pain sensation – a modality that is not critically dependent on PIEZO2 transduction^3^. Here, we identified a distinct subtype of these nociceptors, which responded to cooling and displayed selective tuning to painful hair-pulling. Their preferred stimulus appeared to be mechanical, with the response to cooling relatively modest. The hair-pull pain perception was dependent on Aβ inputs, as shown by the preferential nerve block data and observations in patients with Aβ deafferentation. This is in contrast to pinprick pain, which is reduced but not abolished in patients with Aβ deafferentation^3^. The identification of functional and molecular characteristics of HFNs will enable the exploration of other potential functions of these neurons in both health and disease. For instance, the presence of TRPM8 invites speculation that under atypical conditions, HFNs might become robustly activated by cooling. Notably, chronic cold allodynia in patients with non-freezing cold injury is abolished during a conduction block of myelinated fibers^40^.

HFNs show a high degree of conservation with similar transcriptomic identity, transduction mechanism, morphology, and function between mice and humans. However, conduction speed might be an exception, with mouse hair-follicle nociceptors signaling in the Aδ range^26^, while convergent data from human psychophysics and microneurography suggests Aβ involvement.

We found that hair-pull pain was distinct at both perceptual and behavioral levels, showing an unexpected level of stimulus tuning mediated by a PIEZO2-dependent class of A-fiber mechanonociceptors. Our demonstration that PIEZO2 is critical for HFNs’ response to the hair-pull stimuli is in line with recent data demonstrating that PIEZO2 knockout leads to 50% decrease in the fraction of Aδ-HTMRs responding to high-threshold mechanical stimuli^19^. While there is growing evidence that PIEZO2 is involved in transducing pathological pain following sensitization^14-16,41,42^, it is worth noting that, even though PIEZO2 contributes to some acute pain responses in mice^14^, we demonstrated for the first time that PIEZO2-dependent nociceptors were necessary for perceiving a submodality of acute pain.

Recent insights into human sensory neuron transcriptomics provide a unique opportunity to explore cell classes associated with pain and elucidate the cellular mechanisms underlying painful sensations. However, linking function to transcriptomics is challenging as the transcriptomic data often do not offer an unequivocal prediction. For example, based on transcriptomics, human KIT^+^ neurons are identified as a class of cold nociceptors^5^, mechanonociceptors^4^, or a unique human class of nociceptors with unclear function^6^. However, only after we combined transcriptomics with functional and pharmacological profiling were we able to identify these cells as HFNs with clear functional specialization. Our approach offers a necessary framework for the identification of nociceptor classes that lack clear counterparts in mice and could also be relevant for studying pathologies characterized by robust plastic changes.

We have identified and characterized a distinct form of pain and delineated its cellular and molecular organization in humans. Further, we identified a specialized type of PIEZO2-dependent, fast-conducting nociceptor with a distinct pharmacological profile. This will advance our understanding of the cellular and molecular organization of the pain system and facilitate future studies on the functional roles of these neurons during health and disease.

## Acknowledgements

We acknowledge the Microscopy Core Facility at the Faculty of Medicine and Health Sciences, Linköping University and dr. Vesa Loitto for providing assistance in confocal imaging. We appreciate the help of Dr. Salvador Amezcua with human skin biopsies and Dorota Persson and L. Medling with participant recruitment. We wish to extend our gratitude to Dr. Patrik Ernfors for his advice throughout the project. We are grateful to Dr. Frank Rice for his advice on human biopsies immunostaining. We also want to thank Sandra Donkervoort for help in PIEZO2 deficient syndrome subject recruitment. This work was supported by Swedish Research Council Starting Grant no. 2020-01107 and Knut and Alice Wallenberg Foundation Fellowship (to M.S), Swedish Research Council Project Grant no. 2021-03054 (to S.S.N.), Knut and Alice Wallenberg Project Grant no. 2019.0047 (to H.O), Knut and Alice Wallenberg Clinical Scholar Grant no. 2019.0487 (to H.O.), ALF Grants Region Östergötland (to S.S.N.), Swedish Society of Medicine Project Grant (to S.S.N.), Magnus Bergvalls Stiftelse Research Grant (to S.S.N.), Western Sydney University Funding Scheme (D.A.M.) and Intramural Research Program of the NIH, specifically the NCCIH (to A.T.C.). We are also thankful to other members of the Szczot, Nagi, Olausson, Larsson, and Mahns groups for helpful discussions.

## Author contributions

M.S., S.S.N., and H.O. conceived the study. O.B., E.K., and O.T. conducted and analyzed the psychophysical and reflex experiments in control participants with inputs from S.S.N., and D.A.M. M.B. performed and analyzed murine *in vivo* imaging with assistance from K.T. O.B., S.S.N., and A.G.M. conducted microneurography experiments with assistance from K.K.W.N., and W.M. S.S.N., A.G.M., and D.A.M. conducted the pharmacological mapping experiments. O.B. analyzed all microneurography data. J.L., E.F., and M.S. conducted and analyzed psychophysical testing in individuals with PIEZO2-deficiency syndrome. J.L., E.F., and M.S. conducted and analyzed psychophysical testing in G.L. as well as control participants coupled with testing in PIEZO2-deficiency syndrome individuals. S.S.N., J.C., and O.T. performed psychophysical testing in I.W. D.S., D.B-G, C.B., and A.T.C. coordinated the recruitment of individuals with PIEZO2-deficiency syndrome. H.M. performed and analyzed human immunohistochemistry with input from M.L., and M.B. A.K., and G.G. performed skin mechanics measurements. H.Y. performed and analyzed human fluorescence in situ hybridization experiments. M.B., G.B.C., and F.d.F. performed and analyzed mouse FISH. M.B., and M.L. performed immunofluorescence in mouse tissue. B.R. and I.S. developed quantitative hair-pulling equipment and control software. O.B., and M.B. produced the figures. O.B., M.B., S.S.N., and M.S. wrote the manuscript with input from W.L., D.M., M.L., A.T.C., and H.O.

## MATERIALS AND METHODS

### Human sample

We investigated the mechanisms of hair-pull pain in a series of experiments. First, psychophysical testing for hair-pull pain was performed in six individuals with PIEZO2^DS^ (3 males, 3 females; 12-36 years) and seven control participants (4 males, 3 females; 20-25 years). Individuals with PIEZO2^DS^ had a very similar phenotype based on their clinical presentation and case history as reported previously (Chesler et al., 2016; Szczot et al., 2018). Next, hair-pull pain discrimination, nociceptive reflex, and urge-to-move responses were investigated separately in five (all males; 24-32 years), three (all males; 24-36 years) and 11 control participants (all males; 23-62 years), respectively. Skin deformation measurements were conducted in three control participants (1 female, 2 males; 25-30 years). Next, hair-pull pain coupled with nerve ischemic block was tested in 20 control participants (13 males, 7 females; 21-41 years). In addition, hair-pull pain was tested in two individuals with selective Aβ deafferentation (IW: male, 70 years; GL: female, 64 years). Next, ultrasound-guided microneurography recordings were performed from the radial, dorsal antebrachial, or superficial peroneal nerve of 94 control participants (53 males, 41 females; 19-53 years). Next, human skin biopsies were taken from the distal leg of five control participants (all males, 33-40 years). Finally, human DRG tissues were procured from National Disease Research Interchange (NDRI) comprising six DRGs between T11 to L5 of three human donors (23-61 years).

The exclusion criteria for control participants were neurological or musculoskeletal disorders, skin diseases, diabetes, and pain-relieving or psychoactive medication. All adult participants, as well as guardians of patients with PIEZO^DS^, provided informed consent in writing before the start of the experiment. The studies reported herein were approved by the ethics committee of Linköping University (2016/433-31, 2020-04426, and 2020-04207), National Institutes of Health’s Combined Neuroscience Ethics Committee, Liverpool John Moores University (14/NSP/039), University of Pennsylvania (RLUW1 01), and Western Sydney University (H13204) and complied with the Declaration of Helsinki.

### Psychophysical and reflex measurements

#### Single hair-pull pain task

The participant was comfortably seated in an adjustable chair. Single hair-pull stimuli were delivered using a custom-built handheld hair puller with force-feedback. The hair was secured in place with a small, lightweight clamp, which was attached to a flexible string connected to a mechanical arm. Then, a trained experimenter (O.B.) slowly pulled the hair, and the participant was asked to rate the intensity of pain on a visual analog scale (VAS) ranging from 0 (no pain) to 10 (worst imaginable pain). A short version of the McGill pain questionnaire was used to evaluate the quality of pain. Hair-pull testing was conducted in a similar manner for the Aβ deafferented individual (I.W.). During hair-pull testing in G.L. and PIEZO2^DS^ individuals and matched controls, they were asked to terminate the trial when they perceived pain sensation. If no pain was reported during the trial, we continued pulling the hair until it was plucked out, and collected reports about pain perceived during the trial.

#### Hair-pull pain discrimination task

The experiment consisted of three steps. First, hair-pull pain intensity ratings were matched to a 512-mN pinprick stimulus (QST, MRC Systems GmbH, Heidelberg, Germany). Single hair-pull was tested as described above. Hair-pull and pinprick stimuli were sequentially delivered within a 10-mm radius on the forearm. We observed that a ∼50-mN hair-pull force produced the same pain intensity as a 512-mN pinprick indentation force on a VAS. Then, these matched stimuli were delivered in a random order guided by a computer script. After each trial, the participant was asked whether they detected a hair pull or a pinprick and to rate the pain on a VAS to ensure that pain ratings remained similar for the two stimuli during each discrimination task. Each stimulus was delivered 15 times during this task. The quality of pain was also assessed using the short form of the McGill pain questionnaire, with a further 10 trials per stimulus.

#### Nociceptive reflex

Hair-pull stimuli were delivered using an automated custom-built multi-hair puller made of a speed controlled linear motor with a force sensor. Multiple (∼10-15) hairs were individually threaded through a 3D-printed grid using tweezers and secured in place with 3M™ Transpore™ surgical tape (3M, Solna, Sweden). Four self-adhesive electrodes (Kendall™ H92SG, Cardinal Health, Dublin, US) were attached to the skin overlying the anterior and posterior compartments of the upper arm muscles, and a reference electrode was placed near a bony part of the skin. The muscle activity was recorded (FE238 Octal Bio Amp and PowerLab 16/30, ADInstrument, New Zealand) and plotted in LabChart (v8.1.24, ADInstrument, New Zealand). The latency of the reflex response was calculated from when the voltage (force) from the hair puller deviated from the baseline to when the muscle (EMG) response was initiated. During each trial, the participant was asked to report the intensity of pain on a pain VAS.

#### Urge to move task

Hair-pull were delivered using an automated custom-built pneumatic multi-hair puller. The heat stimuli were delivered using a thermode with a 4.5 cm^2^ stimulation surface and a maximum temperature of 50°C (T09 probe, QST, Strasbourg, France). Initially, the intensity of each stimulus was adjusted to achieve a pain rating of 4 or higher on a pain VAS. Once the stimulus intensities were set, we started the urge to move task. A scale was presented on the screen with the question “How strong was the urge to move?” and “towards” and “away” as endpoints. The two stimulus types were delivered in a randomized sequence with a total of 40 trials, 20 for each modality. Pain ratings were randomly collected during the experiment to ensure that the stimuli remained painful (VAS ≥ 4) throughout the experiment. If the pain rating dropped below 4, the intensity of the stimulus automatically increased until the pain returned to the target range or reached the maximum stimulus range.

#### Nerve ischemic block

An ischemic block was applied to examine the contribution of different afferent fiber classes to the perception of single hair-pull pain. Briefly, a blood pressure cuff was wrapped around the upper arm or above the ankle of the participant and rapidly inflated to 300 mmHg^20^. Then, selective sensory stimuli were applied to gauge the progression of the nerve block, with vibration intensity and detection of cooling and warming taken as measures for Aβ-, Aδ-, and C-fiber function, respectively^21^. Low and high vibration frequencies were delivered using a piezoelectric device (Piezo Tactile Stimulator, Dancer Design, St. Helens, UK), and intensity ratings were collected in triplicate. Thermal stimuli were applied using a contact thermode (Thermal Cutaneous Stimulator TCS II, QST lab, France), with the baseline temperature adjusted to the skin. Cold and warm perception thresholds were assessed by decreasing or increasing the temperature of the thermode at a constant rate (1°C/s) until the participant indicated their perception of cold or warm by pressing a button. Detection thresholds were measured thrice for each modality. After confirming that vibration sense was blocked while temperature was still detectable in the nonpainful range, we evaluated the participants’ ability to detect and rate single hair-pull forces on a pain VAS.

### Microneurography

Single-unit axonal recordings were performed in awake control participants, as described previously (Nagi et al., 2019; Yu et al., 2023). Under real-time ultrasound guidance (LOGIQ P9, GE Healthcare, Chicago, IL, USA), the target nerve was impaled with an insulated tungsten recording electrode (FHC Inc., Bowdoin, ME, USA). An uninsulated reference electrode was inserted adjacent to the recording electrode, just under the skin. A high-impedance preamplifier (MLT185 headstage) was attached to the skin near the recording electrode and used together with a low-noise high-gain amplifier (FE185 Neuro Amp EX, ADInstruments, Oxford, UK). Single units were searched for using soft and rough brushing, pinching, and hair tugging. Mechanical thresholds and receptive field size were determined using Semmes-Weinstein monofilaments (nylon fiber; Aesthesio, Bioseb, Pinellas Park, FL, USA). All recorded afferents were mechanically responsive and divided into subtypes based on established criteria (Nagi et al., 2019; Vallbo et al., 1995; 1999). Then, single hairs within the receptive field of the recorded afferent were pulled using a handheld hair-puller, as described in the psychophysics section. Thermal responsiveness was tested by placing a Peltier probe (zone dimension: 7.4 x 12.2 mm, T09, QST.Lab, Strasbourg, France) onto the receptive field. After recording baseline activity for at least 30 s (with the thermode in contact with the receptive field) at a neutral temperature of 30°C, a series of cooling (down to 0°C) and warming (up to 50°C) stimuli were delivered at 30-s intervals. To test TRPM8 desensitization, 1 mg of menthol was prepared in 10% ethanol and diluted in saline solution. After testing the cooling responsiveness, 0.1 ml of menthol was injected around the receptive field where the unit was located in 3 to 5 injections. Finally, conduction velocity of the recorded afferent was estimated from latency responses to surface electrical or mechanical stimulation of the receptive field (FE180 Stimulus Isolator, ADInstruments, Oxford, UK). Electrically and mechanically evoked spikes were compared on an expanded time scale to confirm they originated from the same unit. Neural activity was sampled at 20 kHz and recorded using the ADInstruments data acquisition system (LabChart software v8.1.24 and PowerLab 16/35 hardware PL3516/P, Oxford, UK), then exported to Spike2 (v10.13, Cambridge Electronic Design Ltd., Cambridge, UK). Recorded action potentials were carefully examined offline on an expanded time scale. Threshold crossing was used to distinguish action potentials from noise with a signal-to-noise ratio of at least 2:1, and spike morphology was confirmed by template matching. Recordings were discarded if multiple units were present or if spike amplitudes were not distinct from the noise, preventing secure action potential identification.

### Skin biopsy, tissue clearing and immunostaining

Healthy human skin biopsies were taken from the distal leg (10 cm above the lateral malleolus) with a 3-mm circular punch device under local anesthesia, according to the US and European guidelines (Lauria et al., 2010). Skin biopsies were fixed in 4% paraformaldehyde (PFA, Polysciences Europe GmbH) at 4°C overnight, cryoprotected in 30% sucrose overnight at 4°C, embedded in optimum cutting temperature compound, and sectioned in cryostat (CM1950, Leica Biosystems Inc) at 100 μm. The free-floating skin sections were stained and cleared with RapiClear 1.47 (Sunjin Lab, Hsinchu City, Taiwan), according to the manufacturer’s protocol. Triple-labelling was performed with combinations of anti-PGP 9.5 (1:1000; ThermoFisher), anti-α-NEFH (1:2000; ThermoFisher) and anti-CGRP (1/250; synaptic system) at 4°C with 5 days of incubation. Then, the immunoreactivities were visualized with the following secondaries antibodies: Alexa 488/568/647 IgG, 1:200; Invitrogen. Photomicrographs were acquired using a spinning disk confocal microscope (Nikon) with a 40x objective.

### Skin Imaging

Biomechanical measurements of skin surface deformation to indentation and hair pulling were assessed using a DIC imaging approach (3D digital image correlation), as previously described (Kao et al., 2021). Indentation and hair pulling were tested thrice per participant. The skin was indented with a 0.6-mm diameter tip, advancing at 5 mm/s with 1 mm increments up to approximately 8 mm or when the participant reached pain threshold. For hair pulling, a single hair was attached to a needle and fed into the indenter tip, adhered with hot wax. The hair was pulled at 5 mm/s with 1 mm increments until it came out. Psychophysical pain ratings were collected for each step within trials.

### Experimental animals

All animal procedures were performed in agreement with Swedish and EU regulations, according to the protocol approved by local ethical commission (DNR214-2021). Following transgenic lines were used: Rosa26-CAG-flox-Stop-GCaMP6f (Ai95D, JAX repository ID 024105, Madisen et al. 2015); Piezo2^fl^ (Szczot et al. 2018), Trpm8:iCre (SMOC repository ID NM-KI-225064), B6.Cg-Calca^tm1.1(cre/EGFP)Rpa^ (JAX repository ID 033168). All animals were bred into C57/Bl6 background.

### AAV injection in mouse pups

Virus was injected intraperitoneally into P0-P3 mouse pups. Animals were briefly anesthetized with Isoflurane/Oxygen mix and injected with 1-2 μl of virus (9 x 10^12^ to 3 x 10^13^ virions/ml) diluted with saline to a total volume of 10 μl. Injection was done using a 0.3 ml insulin syringe with a 30G needle (BD). All viral constructs were provided by Canadian Neurophotonics Platform Viral Vector Core Facility.

### *In vivo* calcium imaging of murine trigeminal ganglia

To gain optical access to the trigeminal ganglion a partial decerebration surgery was performed as previously described ^26^ on 8-12 weeks old mice. *In vivo* calcium imaging was achieved using epifluorescence CSE2300 module of Bergamo II microscope (Thorlabs) with 4x 0.2 NA TL4X-SAP objective. CHROLIS light source (Thorlabs) was used with 475, 525 and 590 nm fluorescence LEDs to image GCaMP6f, tdTomato and mNeptune respectively. Images were acquired at 5 Hz with a PCO Panda 4.2 bi CMOS camera, using MicroManager 2.0.1 software (University of California, San Francisco). In experiments using viral calcium-independent fluorescent labelling the surface of trigeminal ganglion was imaged in red/far red channel and then, without moving the specimen, GCaMP6f imaging was performed.

### Mechanical and thermal stimulation of murine skin

Pulling mouse hair was performed by gluing a single hair to the inside of a 10 μl pipette tip filled with UV curable glue. The end of the pipette tip was cut off and a thin cotton thread was put into the wider end before curing the glue. The thread was then attached to a force sensing load cell mounted on an analog micromanipulator (Narishige) that allowed for slow, controlled increase of mechanical force applied to the hair until extraction. TAL221 load cell (HT Sensor Technology) was connected to a NI USB-6343 digitizer (National Instruments) through a Z-SG analog I/O module (Seneca). Gentle brush was performed with cotton tipped applicators. Prior to thermal stimulation and pinching mouse cheek was shaved and depilated using Nair Sensitive cream (Church & Dwight). Temperature stimuli were applied using a custom-built setup consisting of two water baths with hot and ice-cold water and a peristaltic pump (Thermofisher) delivering water to a copper thermode attached to the skin on the cheek. Trigger pulses generated by NI USB-6343 controlled by FlexLogger software (National Instruments) activated the pump and switched pinch valves, which produced the temperature ramps. Thermal probe was mounted on the copper thermode for temperature readout, which was acquired using CL-100 Temperature Controller (Warner Instruments). Force and temperature readouts synchronized with image acquisition were recorded using FlexLogger.

### Calcium imaging analysis

Out of focus areas of the concatenated image stacks were cropped out and motion artifacts were corrected using a custom Fiji/ImageJ2 script based on the ‘Linear Stack Alignment with SIFT’ plugin that registered individual images to the first image. Activated cells were manually selected as regions of interests in ImageJ. Neuropil subtraction was performed using custom MATLAB script, as described previously (Ghitani et al. 2017). Threshold for neuron activation was set at ΔF/F_0_ ⩾ 8%. Out of all single hair-pull trials (up to 10 per experiment) trials with the lowest force to elicit activation were selected for each neuron. All analyses were done using in-house MATLAB and Python scripts.

### Mouse skin confocal imaging

Homozygous TRPM8:iCre mice were injected with an AAV/PHP.S-CAG-LSL-tdTomato virus at P3 and perfused with 4% PFA at 8 weeks old. Back skin was removed, cut into rectangular pieces, and embedded in OCT medium (CellPath). 80-100 μm sections were cut on a cryostat (Leica), stained with DAPI (Advanced Cell Diagnostics) and imaged on a Spinning Disk confocal microscope (Nikon).

### RNAscope *in situ* hybridization of murine DRG, imaging and quantification

6-10 weeks C57BL/6 mice were perfused with 4% PFA and the DRG tissue has been harvested. The tissue was embedded in OCT (CellPath) and sectioned on a cryostat (Leica). 15 mm sections were mounted on SuperFrost Plus glass slides (ThermoScientific). Multiplex *in situ* hybridization (ISH) was done using the manual RNAscope assay (Advanced Cell Diagnostics) according to supplier’s protocol.

### Quantification of histological data

Overlap of transcriptomic markers in DRG neurons was quantified using Fiji/ImageJ2. Representative images (5 sections from 3 animals) were takes using Spinning Disk microscope (Nikon) and ROIs were drawn using free hand tool. TRPM8^high^ and TRPM8^low^ cell populations were distinguished by a ΔF/F_0_ = 100% threshold of Trpm8 RNAscope signal.

### RNAscope *in situ* hybridization of human DRG, imaging and quantification

OCT embedded freshly dissected human lumbar or thoracic DRG tissues were cryosectioned at 20 μm thickness and mounted on glass slides. The slides were stored in −80 °C to preserve RNA integrity. RNAscope Fluorescent Multiplex Reagent Kit and RNASCOPE probes for the targeted genes (Advanced Cell Diagnostics Inc.) were used for multiplex FISH. RNAscope in situ hybridization was performed in accordance with the manufacturer’s instructions. In brief, fresh frozen hDRG sections were fixed, dehydrated, and treated with protease. The sections were then hybridized with the respective target probe for 2 h at 40°C, followed by two to three rounds of signal amplification. The sections were then mounted under coverslips, sealed with nail polish, and stored in the dark at 4°C until imaged. A Leica SP5 confocal microscope was used to capture images and ImageJ was used for image analysis. In some DRG neurons, accumulation of lipofuscin in part of cells caused strong autofluorescence in all channels. These signals were considered as non-specific background and were excluded from the analysis. The percentage of each cluster across all DRG neurons may be a slightly overestimated due to the underestimation in quantification of total neuronal numbers, because some cells have neither multiple FISH signals nor DAPI (4′,6-diamidino-2-phenylindole) nucleus staining signals.

