## Extended figures for "PIEZO2‐dependent rapid pain system in humans and mice"

### Supplementary material - extended figures

Extended figure 1: Pain qualities in response to pinprick and hair-pull stimulation.
Extended Figure 2: Quantifying skin mechanics for hair-pull and indentation.
Extended Figure 3: Somatosensory testing to monitor nerve block progression.
Extended figure 4: Receptive field locations.
Extended figure 5: Hair-pull pain is not mediated by LTMRs.
Extended figure 6: Hair-pull pain is not mediated by LTMRs or cooling-unresponsive A-HTMRs.
Extended figure 7: Molecular profiling of TRPM8$^{low}$ neurons.
Extended figure 8: TRPM8:iCre line faithfully targeted all TRPM8 expressing neurons.
Extended figure 9: Functional characterization of hair pull responses in transgenic models.

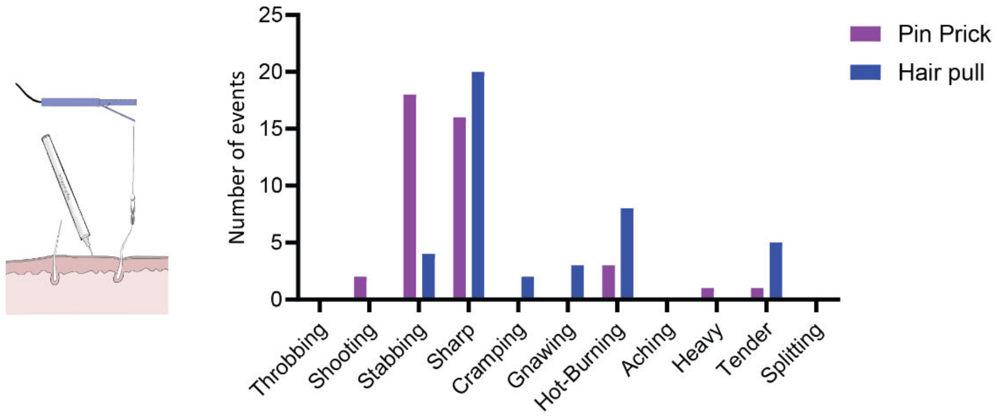


Extended figure 1: Pain qualities in response to pinprick and hair-pull stimulation.

McGill pain quality data in response to hair-pulling and pinprick stimulation at equivalent pain intensities. The data, collected from 5 healthy participants, show the number of occurrences (events) for each pain descriptor, including common descriptors (‘sharp’) and those preferentially chosen for pinprick pain (‘stabbing’) and hair-pull pain (‘hot-burning’ and ‘tender’).

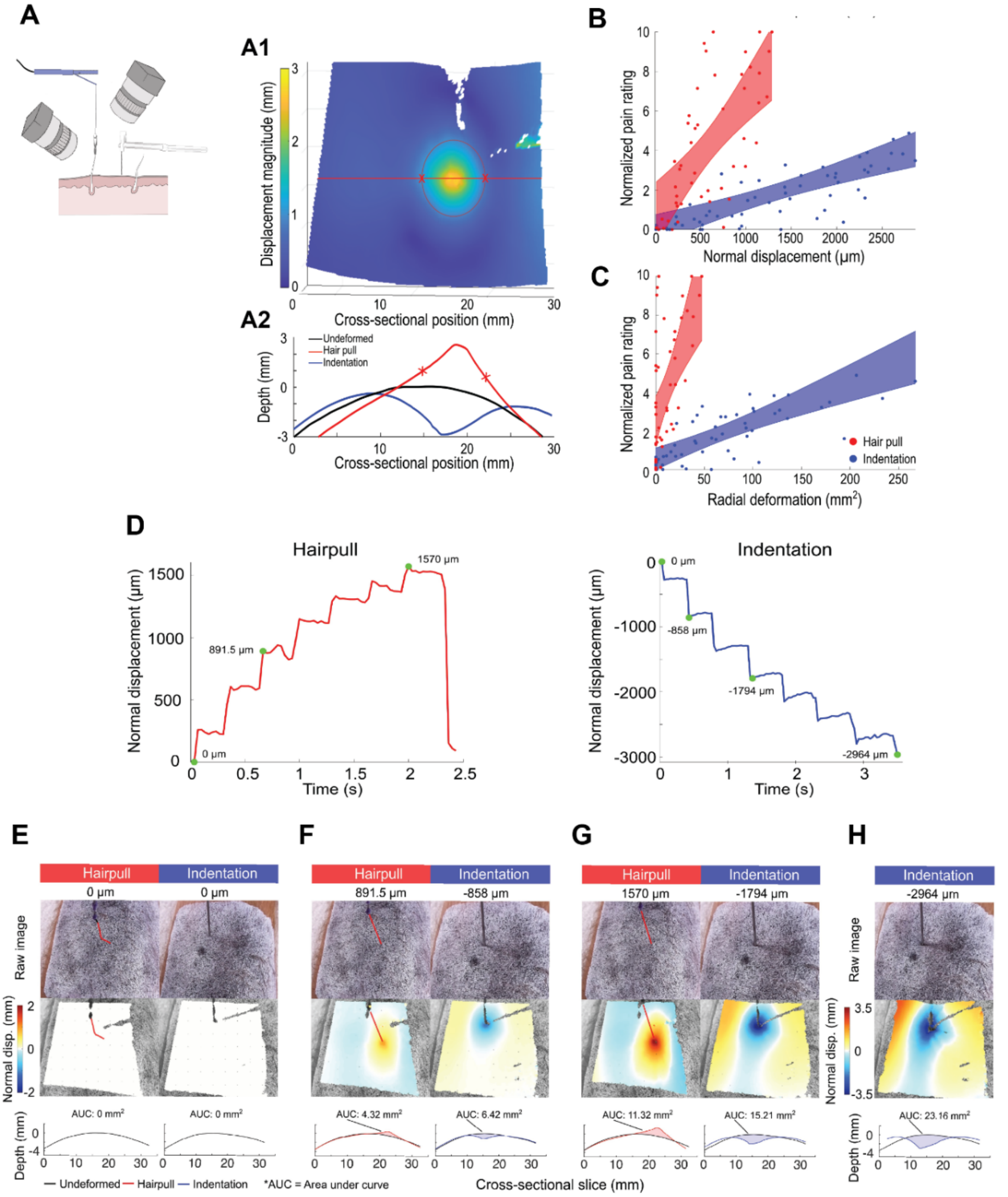


Extended Figure 2: Quantifying skin mechanics for hair-pull and indentation.

A. Comparing states of skin deformation between hair pull and normal indentation using 3D digital image correlation. A1: 3D point cloud showing the change in skin surface displacement from the initial state during hair pull on the forearm (t=0 → t=33 sec). Yellow color depicts greater movement in the x, y, and z directions as compared to purple color depicting no change from the undeformed state. Displacement increases radially inward towards the point of hair pull. The red line marks the central cross-section of the 3D surface, taken at each time point to visualize the normal deformation of the skin surface. From the 3D point cloud, the field of points displacing more than 1 mm are fitted with a 2D ellipse, depicted by the red ring. To quantify lateral deformation, radial deformation is measured as the area of this ellipse. A2: The undeformed cross-section before contact (black, t=0 s), at maximum deformation during the same hair pull trial (red, t=33 s), and at matching depth during a normal indentation trial (blue, t=12.7 s). Similar deformation patterns are exhibited between hair pull and normal indentation, while hair pull extends the skin above the undeformed surface and indentation deforms the skin below the undeformed surface. B. Aggregate raw data of normal displacement for all trials and all participants fit with a linear line and shaded regions depicting the 95% confidence intervals. Higher normalized pain levels are reported at similar levels of normal displacement for hair pull as compared to normal indentation. C. Radial deformation plotted by normalized pain rating showing a steeper slope for hair pull pain as compared to normal indentation pain, despite similar levels of radial deformation, or lateral skin surface movement. Data are represented as average ± SEM. Dots showed the individual trials. D. Raw trace of skin normal displacement over time for one hair pull trial (red) and one indentation trial (blue). Green dots mark the time points visualized in panels E-H below. A maximum normal displacement of 1570 µm is measured during the hair pull trial while a maximum of -2964 µm is measured during indentation, indicating that greater skin deformation can be reached in indentation trails, as compared to hair pull trials, before pain thresholds are reached. E. The raw speckled image, normal displacement, and cross-sectional slice for the same hair pull trial and indentation trial as (E) taken at t = 0 sec, or normal displacement = 0 µm, for the same participant. No difference in initial states was observed and the area under the curve (AUC) was 0 mm2 for both trials. F. At similar steps of normal displacement (hair pull: 891.5 µm, indentation: -858 µm), similar, though inverted, fields of normal displacement and cross-sectional slices were observed. While orange color depicts upward movement for hair pull, and blue color depicts downward movement for indentation, the magnitudes of normal displacement were similar at these two points. Similarly, the cross-sectional slices maintained similar curvature and AUC measurements. G. A further step in normal displacement depicted increasing deformation for both trials, while maintaining similarities in deformation pattern and AUC measurements. H. A final step in indentation, unmatched by hair pull, reinforced how indentation trials can reach greater states of deformation before reaching pain thresholds, as compared to hair pull trials.

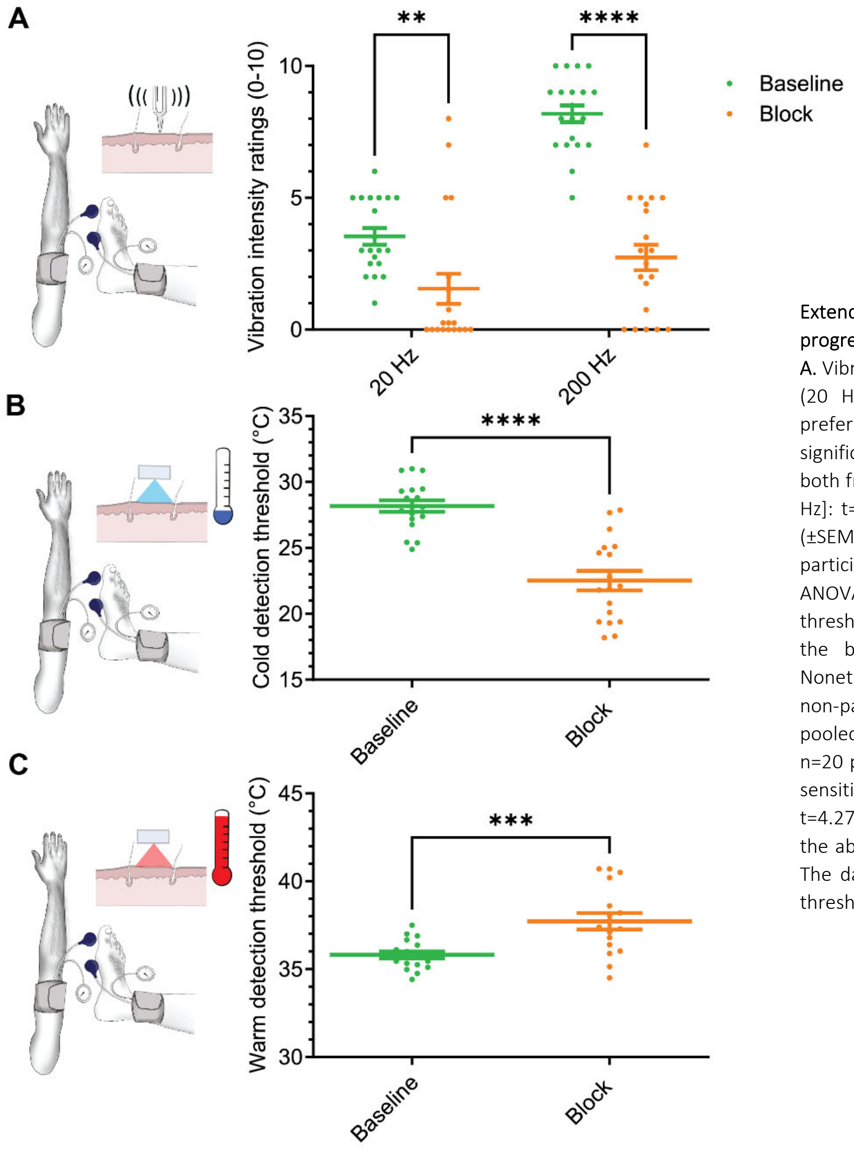


Extended Figure 3: Somatosensory testing to monitor nerve block progression.

**A.** Vibration intensity task. Vibration intensity ratings in response to low (20 Hz) and high (200 Hz) frequencies at 'Baseline' and during preferential ischemic 'Block' of Aβ fibers. The vibration intensity ratings significantly decreased ($F_{(1,19)}$ = 43.85, p<0.0001) during the block for both frequencies (Baseline vs. Block: [20 Hz]: t=3.72, p<0.001 and [200 Hz]: t=10.22, p<0.0001). The data show individual- and pooled-mean (±SEM) vibration intensity ratings (tested in triplicate; n=20 participants). Statistical differences were assessed using one-way ANOVA with Tukey's multiple comparisons test. **B.** Cold detection threshold. Nonpainful cooling sensitivity decreased significantly during the block (Baseline vs. Block: t=6.23, p<0.0001; Paired t-test). Nonetheless, all participants retained the ability to detect cold in the non-painful range (cold pain <10°C). The data show individual- and pooled-mean (±SEM) cold detection thresholds (tested in triplicate; n=20 participants). **C.** Warm detection threshold. Nonpainful warming sensitivity decreased significantly during the block (Baseline vs. Block: t=4.27, p=0.0005; Paired t-test). Nonetheless, all participants retained the ability to detect warm in the non-painful range (heat pain >40°C). The data show individual- and pooled-mean (±SEM) warm detection thresholds (tested in triplicate; n=20 participants).

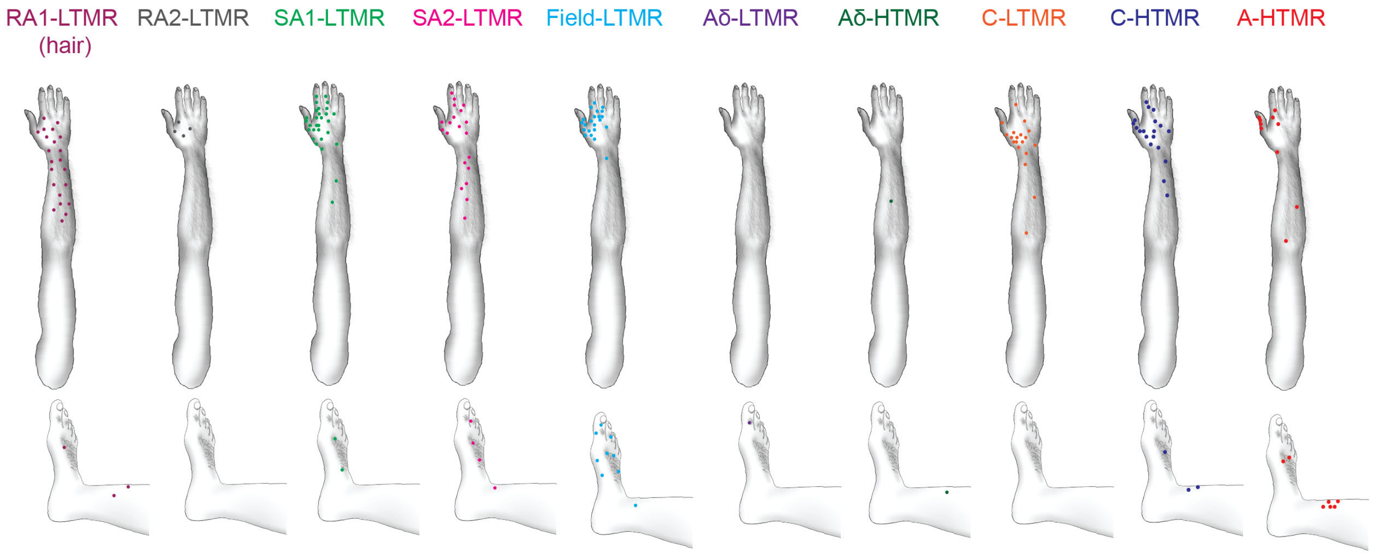


Extended figure 4: Receptive field locations.

Locations of all afferent types from recordings in the peroneal, antebrachial, and radial nerves, with each dot representing an individual afferent (n=180).

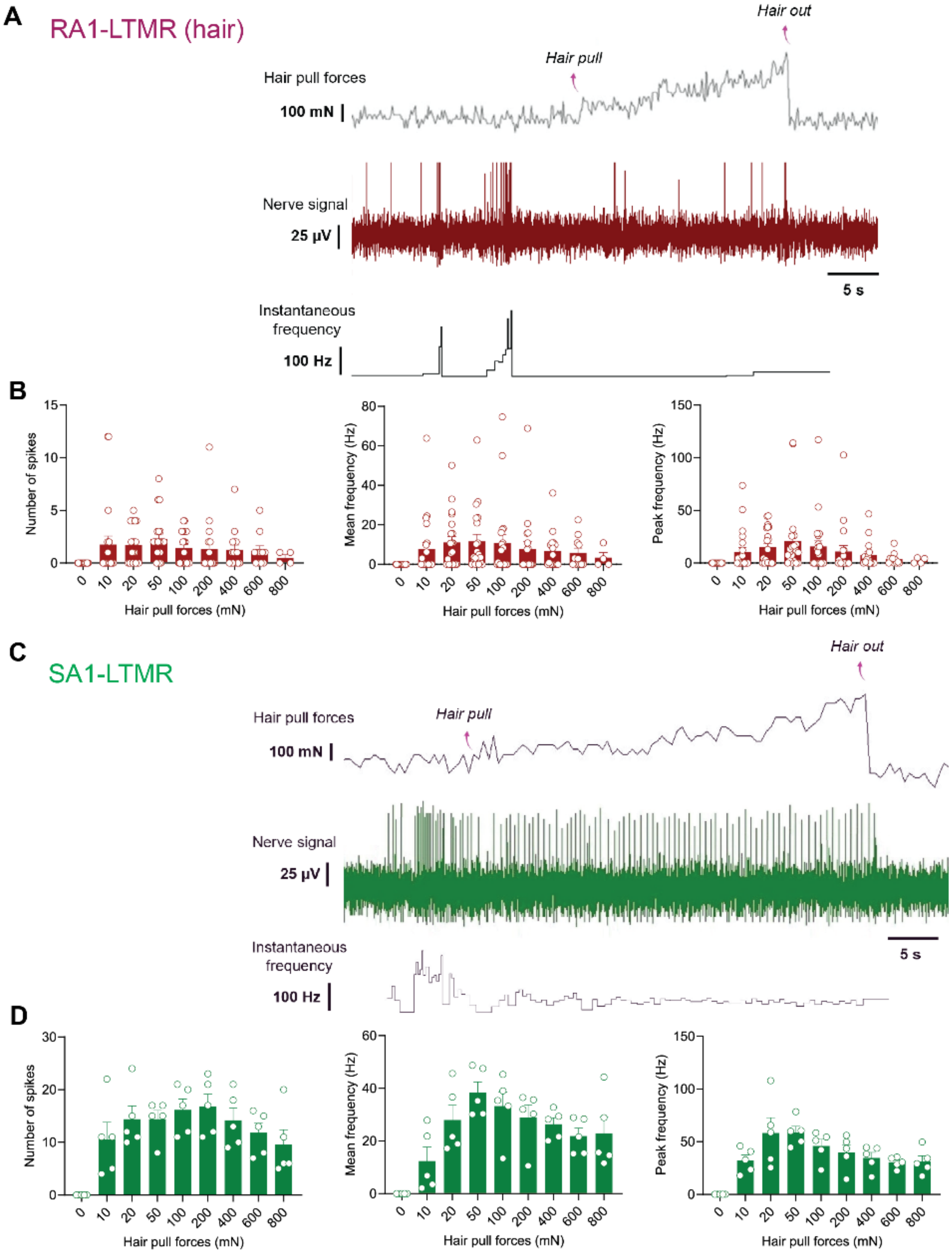


Extended figure 5: Hair-pull pain is not mediated by LTMRs.

A. Representative neural trace showing response of an RA1-LTMR to a single hair-pull. B. Neural discharge (number of spikes and mean and peak frequencies) of RA1-LTMRs (n=13 units). Hair pulling forces had no significant effect on the number of spikes ($F_{(10,168)}$ = 1.861, p=0.0538), mean frequency ($F_{(10,168)}$ = 1.54, p=0.1281), or peak frequency ($F_{(10,168)}$ = 2.13, p=0.0245). The data show individual trials and mean (±SEM) responses of RA1-LTMRs (n=19 trials) to hair pulling at different forces. C. Representative neural trace showing response of an SA1-LTMR to a single hair-pull. D. Neural discharge (number of spikes and mean and peak frequencies) of SA1-LTMRs (n=5 units). Hair pulling forces had no significant effect on the number of spikes ($F_{(10,44)}$ = 8.35, p=0.2855), mean frequency ($F_{(10,44)}$ = 8.44, p=0.3269), or peak frequency ($F_{(10,44)}$ = 9.64, p=0.0321). The data show individual trials and mean (±SEM) responses of SA1-LTMRs to hair pulling at different forces. Statistical differences were assessed using one-way ANOVA with Tukey's multiple comparisons test.

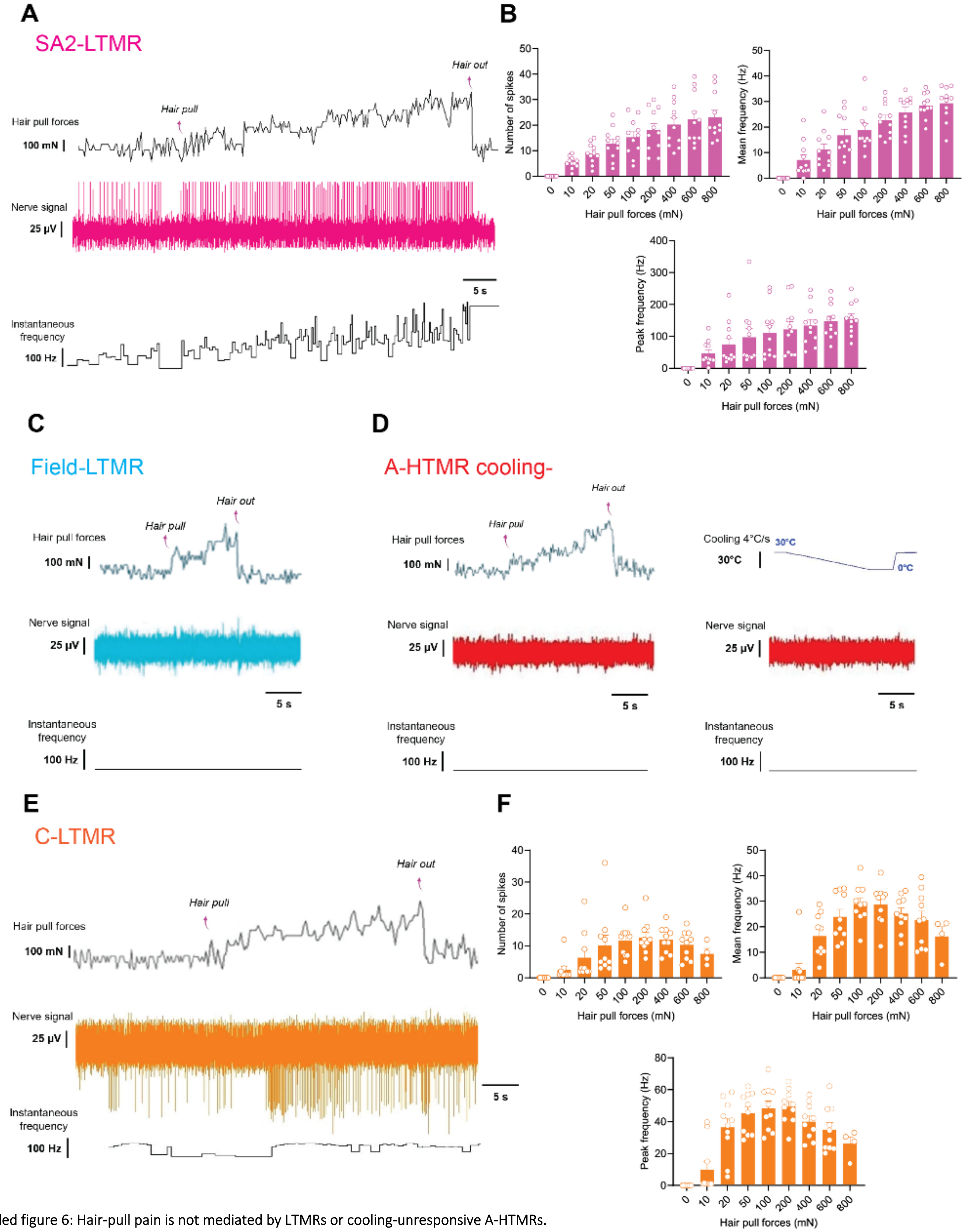


Extended figure 6: Hair-pull pain is not mediated by LTMRs or cooling-unresponsive A-HTMRs.
**A.** Representative neural trace showing response of an SA2-LTMR to a single hair-pull. **B.** Neural discharge (number of spikes and mean and peak frequencies) of SA2-LTMRs (n=7 units). Most SA2s were spontaneously active. Hair pulling forces had a significant effect on the number of spikes ($F_{(10,109)}$ = 18.05, p<0.0001), as well as on the mean ($F_{(10,109)}$ = 35.52, p<0.0001) and peak frequencies ($F_{(10,109)}$ = 9.97, p<0.0001). However, the SA2 responses plateaued during increasing pulling forces. No significant differences were found in the peak frequencies between 800 mN and all forces tested (p>0.05). The data show individual trials and mean (±SEM) responses of SA2-LTMRs (n=11 trials) to hair-pull at different forces. **C.** Representative neural trace of a field-LTMR to a single hair-pull. All field-LTMRs (n=5 units) were unresponsive to hair-pull. **D.** Representative neural traces of a cooling-unresponsive A-HTMR to single hair-pull and cooling. All A-HTMR cooling- (n=7 units) were unresponsive to hair-pull. The mechanical threshold, conduction velocity and cooling response of five Aβ-HTMRs from this sample were previously published in a preprint (Yu et al., 2023). **E.** Representative neural trace showing response of a C-LTMR to a single hair-pull. **F.** Neural discharge (number of spikes and mean and peak frequencies) of C-LTMRs (n=10 units). Hair pulling forces had a significant effect on the number of spikes ($F_{(7,72)}$ = 7.04, p<0.0001), as well as on the mean ($F_{(7,72)}$ = 20.21, p<0.0001) and peak frequencies ($F_{(7,72)}$ = 19.89, p<0.0001). However, no significant effect was seen in the peak frequencies between 800 mN and all forces tested (p>0.05). The data show individual and mean (±SEM) responses of C-LTMRs to hair-pull at different forces. Statistical differences were assessed using one-way ANOVA with Tukey's multiple comparisons test.

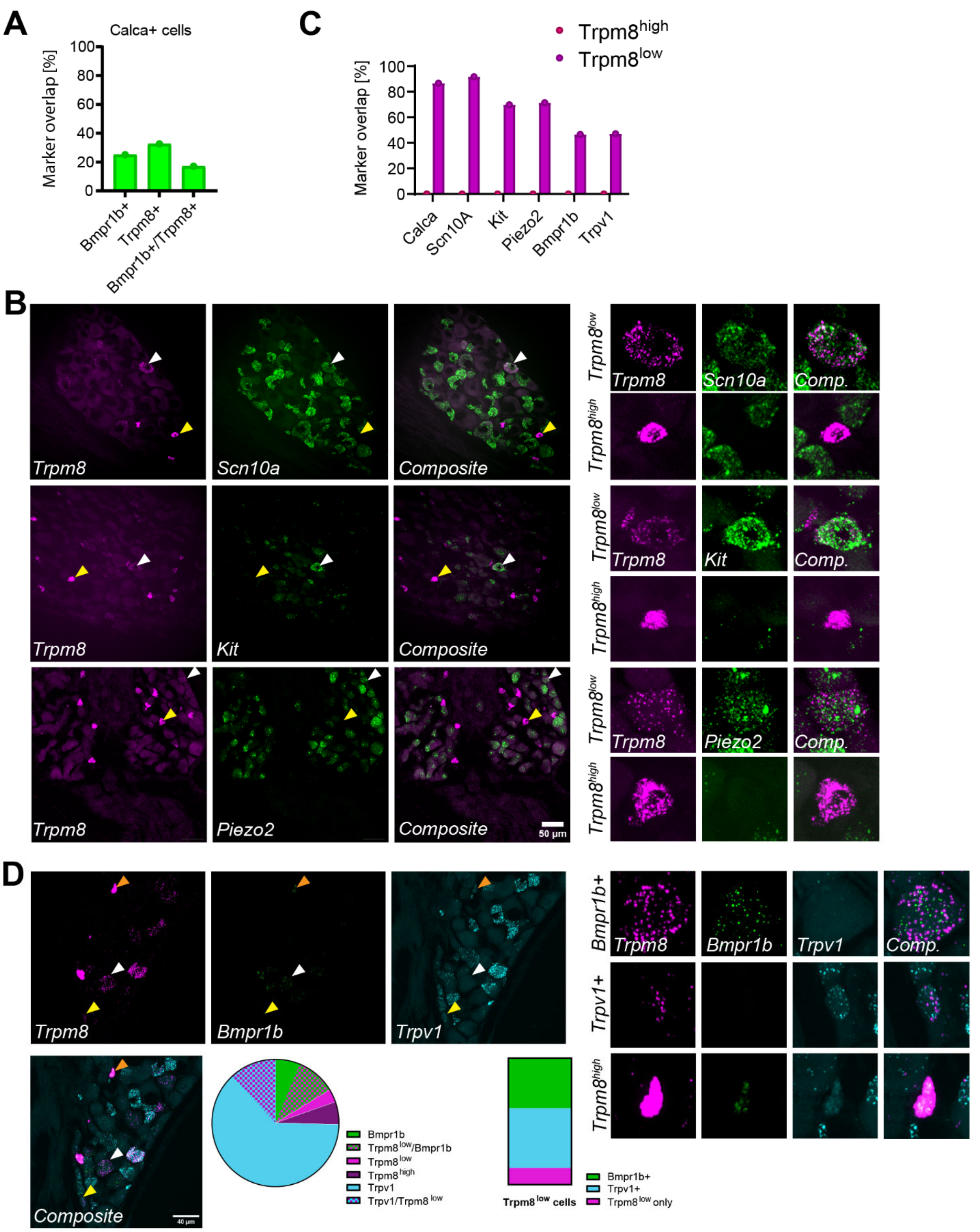


Extended figure 7: Molecular profiling of TRPM8$^{low}$ neurons.

A. Quantification of fluorescence *in situ* hybridization (FISH) of murine DRG data shown in Fig. 4E. B. FISH of murine DRG cells shows expression of hair follicle nociceptor markers in Trpm8$^{low}$ neurons and lack of overlap in Trpm8$^{high}$ cells. C. Quantification of *FISH* shown in D. Trpm8$^{low}$ cells show high overlap with multiple transcriptomic markers expressed by human HFNs, while Trpm8$^{high}$ neurons do not. FISH shows co-expression of TRPV1 and TRPM8 in TRPM8$^{low}$ cells. Pie chart shows distribution of marker expression. Bar chart shows the distribution of genetic markers specifically in TRPM8$^{low}$ cells.

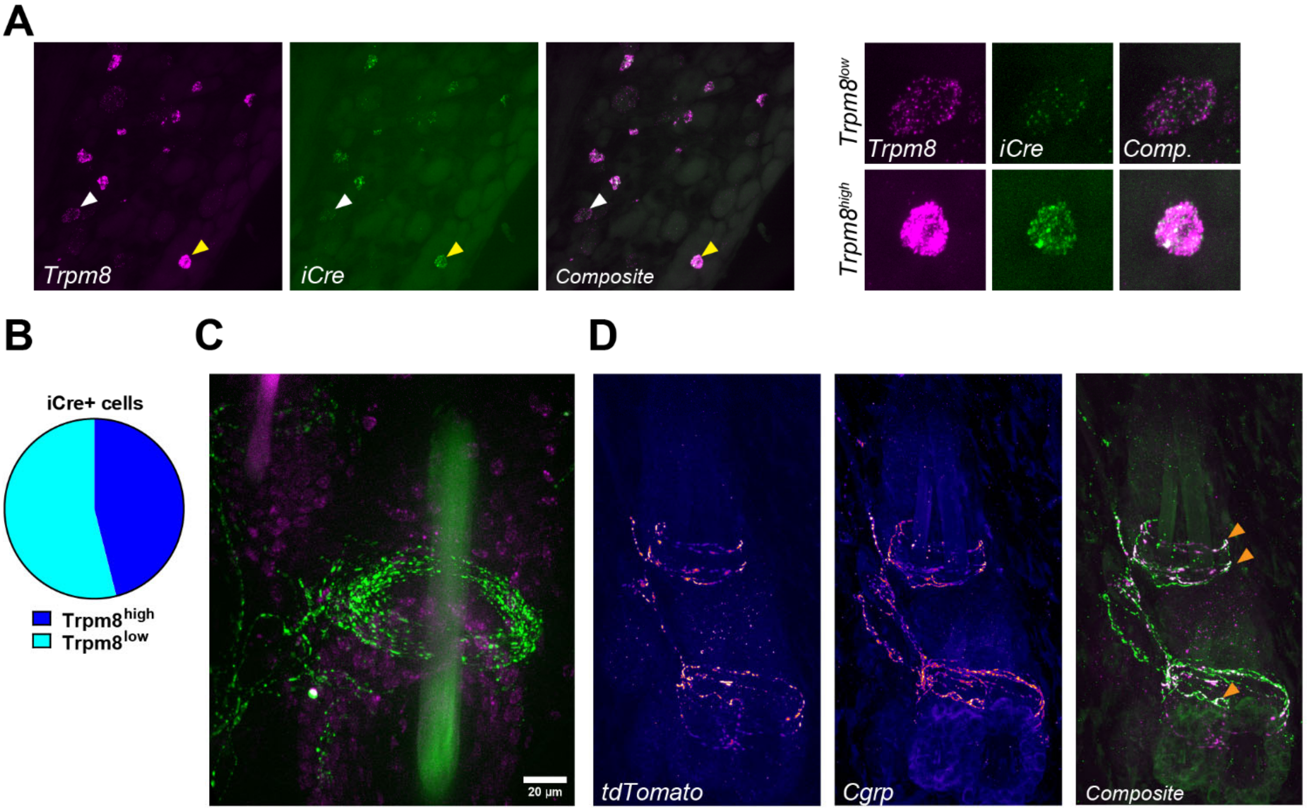


Extended figure 8: TRPM8:iCre line faithfully targeted all TRPM8 expressing neurons.

A. and B. Fluorescence *in situ* hybridization confirms that TRPM8:iCre faithfully targeted both TRPM8$^{low}$ and TRPM8$^{high}$ cells. White and yellow arrows point to Trpm8$^{low}$ and Trpm8$^{high}$ cells respectively, zoomed in on the right. C. Trpm8:iCre efficiently labels circumferential endings in mouse hairy back skin. Trpm8:iCre animals were postnatally injected with cre dependent reporter (AAV/PHP.S-CAG-LSL-tdTomato). tdTomato in green and DAPI in magenta. D. Immunostaining of hairy skin of TRPM8:iCre postnatally labelled with AAV/PHP.S-CAG-LSL-tdTomato mice shows hair follicle-associated nerve endings co-expressing tdTomato and CGRP (green and magenta respectively in the composite panel). Orange arrows point to colocalization hot spots.

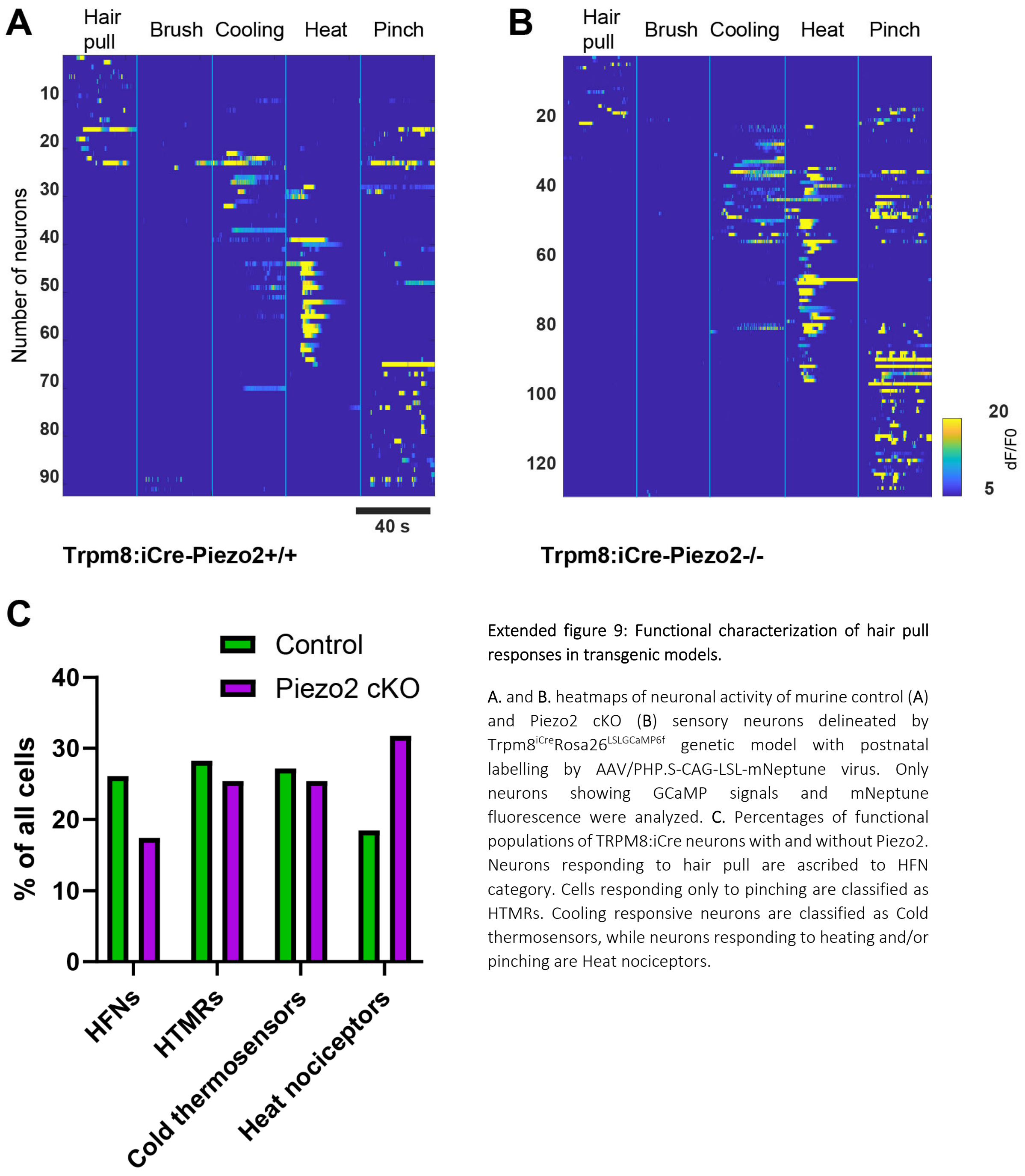


Extended figure 9: Functional characterization of hair pull responses in transgenic models.

A. and B. heatmaps of neuronal activity of murine control (A) and Piezo2 cKO (B) sensory neurons delineated by Trpm8$^{iCre}$Rosa26$^{LSLGCaMP6f}$ genetic model with postnatal labelling by AAV/PHP.S-CAG-LSL-mNeptune virus. Only neurons showing GCaMP signals and mNeptune fluorescence were analyzed. C. Percentages of functional populations of TRPM8:iCre neurons with and without Piezo2. Neurons responding to hair pull are ascribed to HFN category. Cells responding only to pinching are classified as HTMRs. Cooling responsive neurons are classified as Cold thermosensors, while neurons responding to heating and/or pinching are Heat nociceptors.
